# An Orthographic Prediction Error as the basis for efficient Visual Word Recognition

**DOI:** 10.1101/431726

**Authors:** Benjamin Gagl, Jona Sassenhagen, Sophia Haan, Klara Gregorova, Fabio Richlan, Christian J. Fiebach

**Affiliations:** Department of Psychology, Goethe University Frankfurt, Frankfurt/Main, Germany; Center for Individual Development and Adaptive Education of Children at Risk (IDeA), Frankfurt/Main, Germany; Centre for Cognitive Neuroscience, University of Salzburg, Salzburg, Austria.; Brain Imaging Center, Goethe University Frankfurt, Frankfurt/Main, Germany

**Author notes:** **Corresponding Author:** Dr. Benjamin Gagl, Department of Psychology, Theodor W. Adorno-Platz 6, D-60323 Frankfurt/Main.

**Keywords:** Visual word recognition, predictive coding, computational modelling, reaction times, fMRI, EEG, Handwriting.

## Abstract

Most current models assume that the perceptual and cognitive processes of visual word recognition and reading operate upon neuronally coded domain-general low-level visual representations – typically oriented line representations. We here demonstrate, consistent with neurophysiological theories of Bayesian-like predictive neural computations, that prior visual knowledge of words may be utilized to ‘explain away’ redundant and highly expected parts of the visual percept. Subsequent processing stages, accordingly, operate upon an optimized representation of the visual input, the *orthographic prediction error*, highlighting only the visual information relevant for word identification. We show that this optimized representation is related to orthographic word characteristics, accounts for word recognition behavior, and is processed early in the visual processing stream, i.e., in V4 and before 200 ms after word-onset. Based on these findings, we propose that prior visual-orthographic knowledge is used to optimize the representation of visually presented words, which in turn allows for highly efficient reading processes.

## Introduction

Written language – script – developed over the last ∼8,000 years in many different variants (Haarmann, 2007). It is a symbolic representation of meaning, based on the combination of simple high contrast visual features (oriented lines) that our brains translate efficiently into linguistically meaningful units. Cognitive models of reading specify the perceptual and cognitive processes involved in activating orthographic, phonological, and lexico-semantic representations of perceived words from such low-level visual-perceptual features (for a review see Norris, 2013). While some models – consistent with other domains of perception (e.g., Riesenhuber & Poggio, 1999 for object recognition) – indeed assume oriented line representations as the lowest-level visual feature involved in visual word recognition (e.g., Coltheart, Rastle, Perry, Langdon, & Ziegler, 2001; Davis, 2010; Dehaene, Cohen, Sigman, & Vinckier, 2005; McClelland & Rumelhart, 1981; Perry, Ziegler, & Zorzi, 2007; Whitney & Cornelissen, 2008), other cognitive models use as starting point a more integrated, domain-specific representation, i.e., letters (Engbert, Nuthmann, Richter, & Kliegl, 2005; Reichle, Rayner, & Pollatsek, 2003; Sibley, Kello, Plaut, & Elman, 2008). Interestingly, this does not take into account findings from vision neuroscience indicating that already the neuronal representation of ‘low level’ visual feature like an oriented line is an abstraction of the visual input: For example, the well-established phenomenon of end-stopping describes that an oriented line (i.e., the frequently-assumed low-level input into the visual word recognition system) is not represented in the brain by many neurons with receptive fields along the length of the line, but by only two neurons that have their receptive fields at the beginning and end of the line (Bolz & Gilbert, 1986; Hubel & Livingstone, 1987; Hubel & Wiesel, 1965). While preserving the representation of line length and angle, this neuronal representation is more efficient (i.e., simplified two neurons vs. two plus all neurons along the line) by several orders of magnitude. Given these results, we hypothesized that early perceptual stages of visual word recognition should also operate upon optimized representations of the visual-orthographic input – which is so far not accounted for by any of the established models of reading and visual word recognition.

To provide a computationally explicit account for explaining end-stopping, Rao and Ballard (Rao & Ballard, 1999) successfully adapted the computational principles of *predictive coding* (Srinivasan, Laughlin, & Dubs, 1982). Predictive coding postulates that perceived regularities in the world are used to build up internal models of the (hidden) causes of sensory events, and that these internal predictions are imprinted in a top-down manner upon the hierarchically lower sensory systems, thereby increasing processing efficiency by inhibiting the processing of correctly predicted input (Friston, 2005; Rao & Ballard, 1999). When sensory input violates these expectations or is not fully predicted, a *prediction error signal* is generated and propagated up the cortical processing hierarchy in a bottom-up fashion (e.g., Todorovic, van Ede, Maris, & de Lange, 2011), where it is used for model updating and thus serves to optimize future predictions (Clark, 2013; Rao & Ballard, 1999). In the case of line representations and end-stopping, neurons with receptive fields at the beginning and end of the line fire and this information is propagated to higher areas where they activate abstract line representations, which in turn in a recursive, top-down manner ‘predict away’ the activity of the receptive fields between the two endpoints of the line (Rao & Ballard, 1999). Predictive coding has by now received support in many domains of perceptual neuroscience, from retinal coding (Srinivasan et al., 1982), auditory (Todorovic et al., 2011; Wacongne, Changeux, & Dehaene, 2012) and speech perception (Arnal, Wyart, & Giraud, 2011; Gagnepain, Henson, & Davis, 2012) to object (Kersten, Mamassian, & Yuille, 2004) and face recognition (Schwiedrzik & Freiwald, 2017), indicating that this framework is likely a generalized computational principle of the brain.

Most readers can process written language at a remarkably high speed. We reasoned that the high efficiency of visual-orthographic processing necessary for efficient reading makes it likely that the visual system also optimizes the ‘low-level’ perceptual representations used for orthographic processing during reading. Inspired by the wide applicability of the principles of predictive coding (see the previous paragraph), the present model-based study explores whether computational principles of predictive coding may contribute to the optimization of neuronal signals at early cortical stages of the perception of written words. In an influential theoretical paper, Price and Devlin (Price & Devlin, 2011) have proposed that principles of predictive coding may be involved in visual word recognition. Their ‘Interactive Account’ model focuses explicitly on ‘intermediate-level’ stages of visual word processing that are attributed to the left ventral occipito-temporal cortex (lvOT; often also referred to as ‘visual word form area’; e.g., Dehaene & Cohen, 2011; Dehaene et al., 2005). The Interactive Account model postulates that at the level of lvOT, visual-perceptual information that is propagated bottom-up from early visual to higher areas when reading a string of letters is integrated with phonological and semantic information fed to lvOT from higher cortical areas in a top-down manner. Empirical support for this proposal comes, for example, from a study by Kherif et al. (2011) demonstrating semantic priming effects between words and pictures of objects in the lvOT.

However, predictive coding as a general model of cortical processing should, in principle, not be restricted to a specific level of processing, but rather apply to all sensory-perceptual stages of the reading process – including also reading-related visual processes in ‘lower level’ perceptual areas (i.e., that take place before the integrative processes attributed to the vOT/visual word form area in the Interactive Account model; see, e.g., also Fig. 2a of Price & Devlin, 2011). We thus hypothesized here that ‘higher level’ linguistic expectations – either in the form of contextual constraint from preceding input or in the form of our knowledge of the orthography of a language – should be imprinted upon the early stages of visual processing, thereby ‘optimizing’ pre-lexical perceptual processing stages that are typically associated with brain processes located posterior to the visual word form area (y-coordinates < − 60 according to Lerma-Usabiaga, Carreiras, & Paz-Alonso, 2018) and temporally earlier than 250 ms (Grainger & Holcomb, 2009).

A critical indication that one might implement a prediction error based on feature-configurations of letters or words (i.e., our orthographic knowledge) is that these linguistic units contain highly redundant visual characteristics (Changizi, Zhang, Ye, & Shimojo, 2006). For example, vertical lines often occurring at the same position (e.g., the left vertical line in E, R, N, P, B, D, F, H, K, L, M) or letters often positioned at the same location in a word (e.g., *s* or *y* as final letters in English). As such, redundancies contribute very little to the identification of letters and words so that removing the visual-orthographic input for the redundant part of the percept is a plausible strategy of our brain to reduce the amount of to-be-processed visual signal – and thereby increasing the efficiency of the neuronal code that underlies visual word recognition. We accordingly propose that following the principles of predictive coding, the visual-orthographic input signal is ‘optimized’ based on our knowledge and expectations about the redundancies of the respective orthography. In other words, we propose that our orthographic knowledge of the language is used to ‘predict away’ the uninformative part of visual input during reading. As a result, the subsequent stages of visual word recognition (as described in several models of reading; see above and Norris, 2013 for review) can proceed on the basis of an optimized representation of the input. As this optimized signal highlights the unexpected (and thus more informative) part of the stimulus, we termed it the *orthographic prediction error* (oPE), in line with the concept of prediction error as used in the predictive coding framework (e.g., Rao & Ballard, 1999). In the following, we describe one possible, computationally explicit implementation of this proposal, which we refer to as the Prediction Error Model of Reading (PEMoR). Following this, we report a series of quantitative evaluations of this model using lexicon-based, behavioral, EEG, and functional MRI data. To compare our prediction error-based model to word recognition without top-down predictions and prediction errors, we also conducted most of the reported analyses for a ‘full’ low-level representation of the perceived stimuli (based on all pixels of the stimulus image). As a result, for most model evaluations, we compare two parameters, one reflecting strictly bottom-up visual processing without a prediction-based optimization step and one based on a top-down, prediction-based optimization of the sensory representation of the perceived word.

### The Prediction Error Model of Reading

The Prediction Error Model of Reading (PEMoR) postulates that our brain identifies words not on the basis of the full physical input into the visual cortex that is contained in a string of letters but rather based on an optimized (and thus more efficient) neuronal code representing only the informative part of the percept (while redundant and expected signals are canceled out Rao & Ballard, 1999). In the predictive coding framework, this non-redundant portion of a stimulus is formalized as a prediction error; we apply this principle to visual word recognition, and propose that internal (i.e., knowledge- or context-dependent) visual-orthographic expectations are subtracted from the sensory input, so that further processing stages operate upon an *orthographic prediction error* (oPE) signal (Fig. 1a). In this first implementation of the PEMoR, the prediction error signals will be computed at the level of the entire word, as words are the most salient visual units when reading Latin script (because they are separated by spaces). Also, it is known from eye tracking research that critical information about a word (like its length) is available before the word is fixated, due to parafoveal pre-processing (Gagl, Hawelka, Richlan, Schuster, & Hutzler, 2014; Rayner, 1975), so that enough information is available to generate specific predictions about the upcoming input on the fly. However, we would like to point out that prediction errors can also be computed based on other levels of representation, i.e., on the level of letters so that alternative or even complementary implementations can be tested within the PEMoR framework. It is commonly believed that higher-level linguistic representations can initiate specific expectations about upcoming words (DeLong, Urbach, & Kutas, 2005; Kliegl, Nuthmann, & Engbert, 2006; Nieuwland et al., 2018; Price & Devlin, 2011) – e.g., about the class (like noun or verb) and meaning of the next word in a sentence like *“The scientists made an unexpected … (discovery)”*. The fundamental difference between these psycholinguistic assumptions about semantic and syntactic predictions and the proposed *visual-orthographic prediction* in PEMoR is that we postulate predictive processes already at much earlier perceptual stages of visual word recognition.

**Figure 1.**
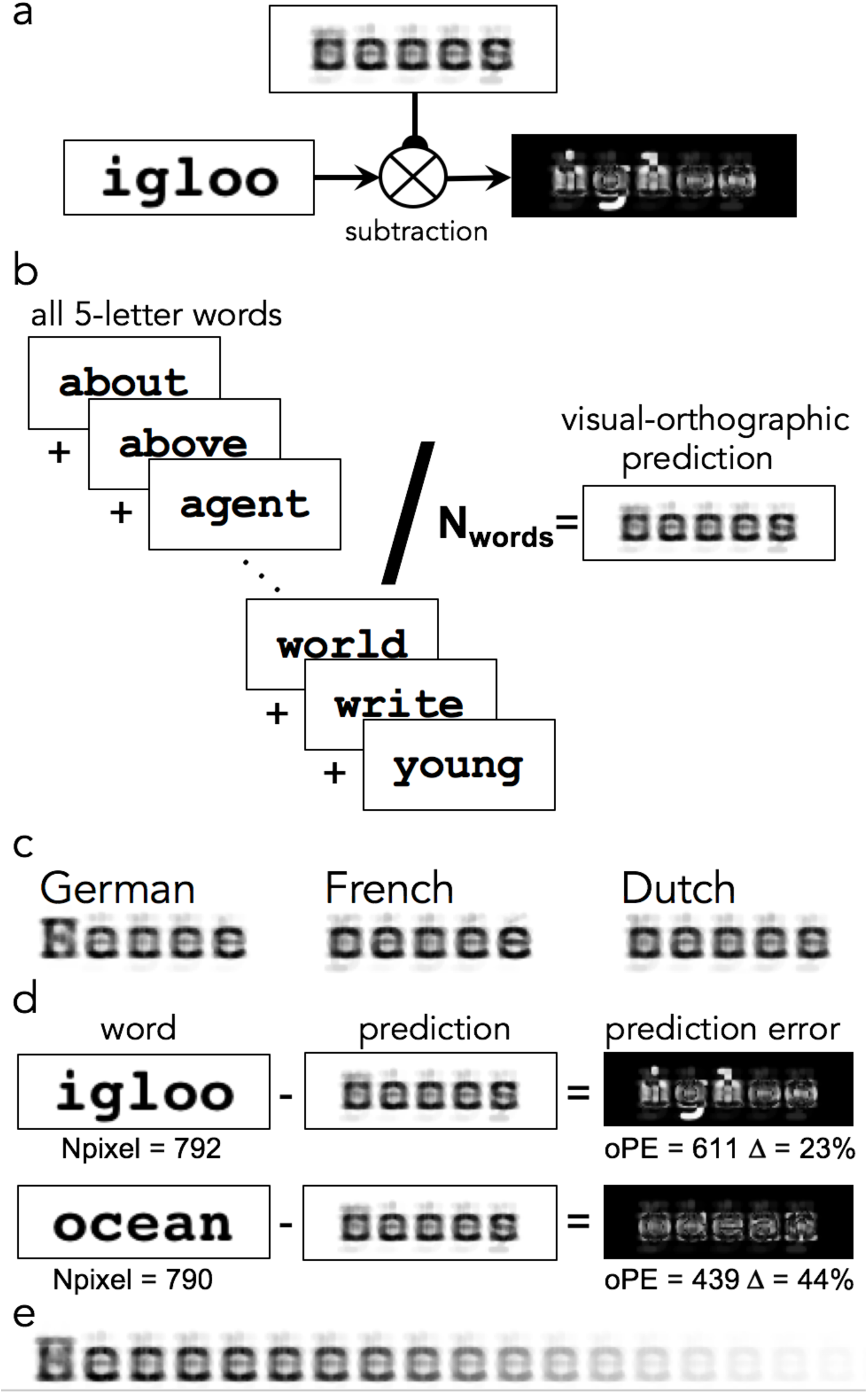
Prediction error model of reading (PEMoR). (a) The PEMoR assumes that during word recognition, redundant visual signals are ‘explained away’, thereby highlighting the informative aspects of the percept. Subsequent stages of word recognition and linguistic processing (i.e., accessing abstract letter and word representations), thus, operate upon an optimized input representation. This assumption is here tested for single-word reading, i.e., independent of context, by subtracting a ‘visual-orthographic prediction’ from the input. (b) The knowledge-based visual-orthographic prediction is implemented as a pixel-by-pixel mean across image representations of all known words (here approximated by all words in a psycholinguistic database; only five letters words, as in most experiments reported here; but see panel (e) of the present figure for predictions including different word lengths and Supplemental Figure S1b). The resulting visual-orthographic prediction, shown on the right, contains the most redundant visual information across all words. (c) Across multiple languages, these predictions are very similar, with the exception of the upper-case initial letter that is visible in the German prediction (because experiments in German involved only nouns and the German orthography requires an upper-case initial letter). (d) The orthographic prediction error (oPE) is estimated, for each word, by a pixel-by-pixel subtraction of the orthographic prediction from the input word (based on their image representations; see Methods for details). While the two example words have similar numbers of pixels, subtracting the orthographic prediction results in substantially different residual (i.e., oPE) images. The values underneath the prediction error images represent a quantitative estimate of the orthographic prediction error, the sum of the gray values of all pixels per image, and show that the amount of signal reduction (Δ) can differ strongly between words. (e) Letter-length unspecific prediction for German, based on ∼190.000 words.

Here, we quantitatively test the assumptions of the PEMoR for the most frequently investigated paradigm in reading research, single word recognition. In the absence of sentence context, the redundant visual information (i.e., the *visual-orthographic prediction* or, in Bayesian terms, the *prior*) is a function of our orthographic knowledge of words. We approximate this prior knowledge quantitatively as the pixel-by-pixel mean over image representations of all words derived from a psycholinguistic database (Brysbaert et al., 2011; see Fig. 1b and Methods). Interestingly, the resulting visual-orthographic predictions look similar across different languages sharing the same writing system (compare Fig. 1c) and are highly intercorrelated (i.e., correlations based on the individual gray values of the pixels from the prediction image; *r* ranging from .95 to .99). Crucially, the high correlations show the robustness of the proposed prediction error representation against such visually salient differences as the capitalization of the first letter in one of the three languages (German). We suggest that these high correlations support the assumption of prediction and prediction error-based word recognition of an abstract principle of word recognition.

We estimate the orthographic prediction error as a pixel-by-pixel subtraction of this visual-orthographic prediction (or prior in Bayesian terms) from each perceived word (Fig. 1d). This step of ‘predicting away’ the redundant part of written words reduces the amount of to-be-processed signals by up to 51% (on average 33%, 37%, and 31% for our English, French, and German datasets, respectively; see Methods, Formula 4), thereby optimizing the visual input signal in the sense of highlighting only its informative parts (Fig. 1d). According to the PEMoR, the resulting orthographic prediction error is a critical pre-lexical stage of word identification, representing (at least part of) the access code that our brain uses to activate word meaning.

We test this model by calculating for each stimulus item a numeric prediction error (oPE) value. This value, i.e., the pre-stimulus sum oPE, equates the sum of all grayscale values from the prediction error image resulting from the subtraction computation of the PEMoR. While this per-item value does not take into account the spatial layout of the stimulus item, it represents an estimate of the amount of neuronal activation needed to represent the specific stimulus. Importantly, representing the oPE as a single value allows us to compare it directly to other typical word characteristics that are closely tied to different psychological models, like word frequency (Brysbaert et al., 2011) or orthographic familiarity (Coltheart, Davelaar, Jonasson, & Besner, 1977; Yarkoni, Balota, & Yap, 2008).

Following the comparison of computational models of reading described by Norris (2013), we consider the PEMoR to be a computational/mathematical model (as opposed to symbolic and interactive activation models), that describes a circumscribed component process of word recognition, i.e., optimization of the perceptual-orthographic representations (as opposed to modeling tasks like lexical decision or reading aloud), and that is based on a rather extensive lexicon. While current models of visual word recognition (e.g., Coltheart et al., 2001) typically act on abstracted representations of the word stimuli (i.e., at the level of lines or letters), the PEMoR operates on the level of the pixels that make up the visual-orthographic stimulus, reflecting its explicit focus on relatively ‘low level’ perceptual-orthographic processing that operate prior to even the earliest processing stages of most other models of visual word recognition. Lastly, the PEMoR differs from previous models in that it is (i) explicitly rooted in a neurobiological model of cortical function (i.e., the predictive coding framework), (ii) has clear assumptions about the neuroanatomical localization of the respective process in the brain, and (iii) in that it is accordingly also directly evaluated against neurophysiological data.

In the following, we provide empirical support for this model by demonstrating that our orthographic prediction error (i) is correlated with orthographic familiarity of words measured as a property of lexicon statistics, (ii) accounts for lexical decision times in three languages, (iii) is represented in occipital brain regions, and (iv) electrophysiological signals from 150-250 ms after word onset. As there exists – to the best of our knowledge – no generally accepted null model against which to compare the PEMoR (or any other computational model of word recognition) and given that no other model of visual word recognition specifies processes at the level of the first cortical stages of visual processing, we also quantified the visual signals contained in each stimulus item prior to prediction-based optimization, by calculating the sum of all pixels in each original stimulus image (see left column of Figure 1d). We used this pixel count parameter as an estimate of the full bottom-up signal that would have to be processed in the absence of prediction-/top down-based optimization of the percept. For most empirical model evaluations reported in the following, we thus compare the performance of the orthographic prediction error against the pixel count parameter.

## Materials and Methods

### Implementation of the Prediction error model of reading

The estimation of the orthographic prediction error, as assumed in PEMoR, was implemented by image-based computations. Using the *EBImage* package in *R* (Pau, Fuchs, Sklyar, Boutros, & Huber, 2010), letter-strings were transformed into grayscale images (size for, e.g., 5-letter words: 140×40 pixels) that can be represented by a 2-dimensional matrix in which white is represented as 1, black as 0, and gray as intermediate values. This matrix representation allows an easy implementation of the subtraction computation presented in Fig. 1a, i.e.,

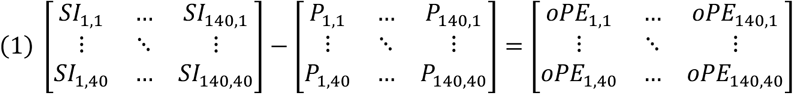

where *SI_x,y_* indicates the sensory input at each pixel. *P_x,y_* reflects the prediction matrix, which is in the present study calculated as an average across the *SI_x,y_* of all words (or a subset thereof) in a lexical database such as the example shown in Fig. 1b based on 5,896 nouns of five letters length from the English SUBTLEX database (Heuven, Mandera, Keuleers, & Brysbaert, 2014):

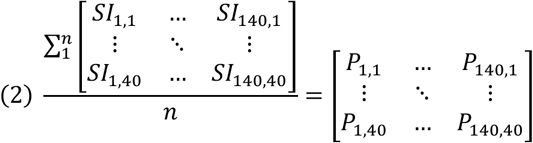

The PEMoR model postulates that during word processing, *SI* is reduced by the prediction matrix *P*, resulting for each stimulus, in an orthographic prediction error matrix (*oPE*), as shown above in formula (1). The resulting orthographic prediction error is, therefore, black (i.e., value 0) at pixels where the prediction was perfect and gray to white (i.e., values > 0) where the visual signal was not predicted perfectly. As the last step, a numeric value for the orthographic prediction error of each stimulus, *oPE_sum_*, was determined by summing all values of its prediction error matrix. This numeric representation of the prediction error is used as a parameter for all empirical evaluations.

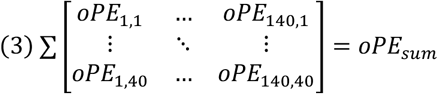

The amount of signal reduction (*S_reduced_*) achieved by this predictive computation can then be calculated by relating the numeric representation of the prediction error to an analogous numeric representation of the respective word’s input image, *SI_sum_*:

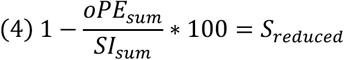

### Participants

35, 54, 39, 31, and 38 healthy volunteers (age from 18 to 39) participated in the two lexical decision studies (i.e., deciding behaviorally if a letter string is a word or not), an fMRI study, an EEG study, and a handwriting experiment, respectively, conducted in German. All had normal reading speed (reading scores above 20th percentile estimated by a standardized screening; unpublished adult version of Mayringer & Wimmer, 2016), reported absence of speech difficulties, had no history of neurological diseases, and normal or corrected-to-normal vision. Participants gave written informed consent and received student credit or financial compensation (10€/h) as an incentive for participating. The research was approved by the ethics board of the University of Salzburg (EK-GZ: 20/2014; fMRI study) and Goethe University Frankfurt (#2015-229; EEG study, lexical decision studies). Behavioral results from 78 English and 975 French readers were obtained from publicly available datasets of mega-studies (for details see Ferrand et al., 2010; Keuleers, Lacey, Rastle, & Brysbaert, 2012).

### Materials, experimental procedures, and statistical analyses

#### Lexicon-based Characterization of the Orthographic Prediction Error

We calculated the number of pixels per word, the orthographic prediction error as described above, and established word characteristics, i.e., the Orthographic Levenshtein distance (Yarkoni et al., 2008) and word frequency, for all five-letter words of each language, i.e., 3,110 German words (Brysbaert et al., 2011), 5,896 English words (Heuven et al., 2014), 5,638 French words (New, Pallier, Brysbaert, & Ferrand, 2004), and 4,418 Dutch words (Keuleers, Brysbaert, & New, 2010). For the German, we additionally estimated a more comprehensive set of orthographic word characteristics, including bi-, tri-, and quadirgram-frequencies (i.e., occurrences of 2, 3, 4 letter combinations), and Coltheart’s N (Coltheart et al., 1977); see Fig. 2b. Orthographic Levenshtein distance and Coltheart’s N were estimated with the *vwr* Package in R (Keuleers, 2013).

**Figure 2.**
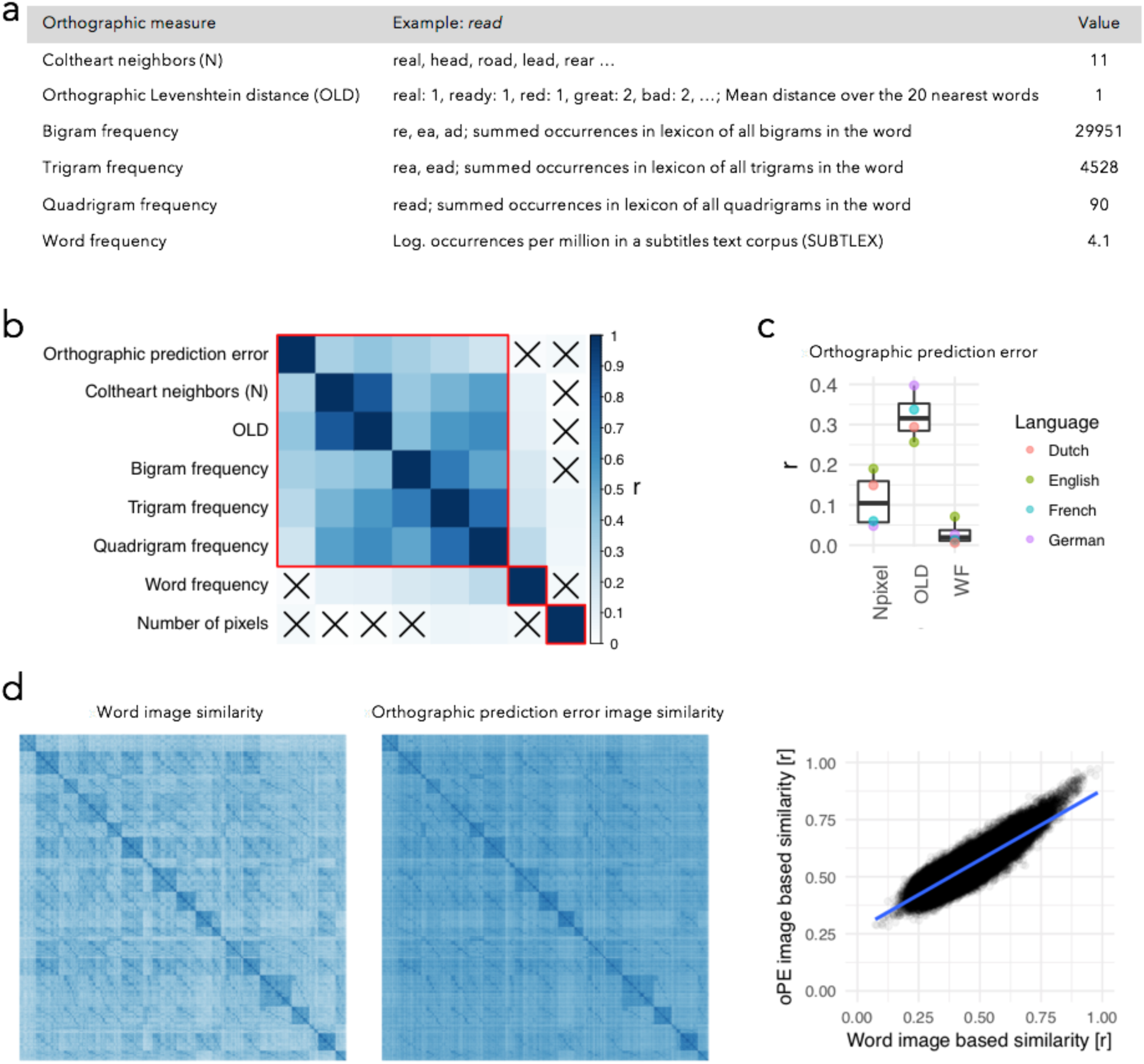
Comparison of orthographic prediction error to established lexicon-based word characteristics. (a) Overview of established word characteristics, exemplified for the word ‘read’: Coltheart’s neighborhood size (Coltheart N; Coltheart et al., 1977), orthographic Levenshtein distance (OLD20; Yarkoni et al., 2008), sub-lexical frequency measures (bi-, tri-, and quadrigram frequencies, i.e., number of occurrences of two, three, and four-letter combinations from the target word, in the lexicon), and word frequency as calculated from established linguistic corpora (see Methods for details). (b) Clustered correlation matrix between the orthographic prediction error, the number of pixels per original image, which represents an estimate of the pure amount of physical bottom-up input in the present study, and the described word characteristics (cf. panel a for explanations), applied to a set of 3,110 German nouns. Red rectangles mark clusters (obtained from a standard hierarchical clustering algorithm using the dendrogram) and black crosses mark non-significant correlations (tested at p < .05; Bonferroni corrected to p < .00179). Number of pixels refers to the original stimulus item and is used as a simplified model of the full bottom-up physical input (Baseline model; see text). (c) Correlations between the orthographic prediction error and number of pixels per word (Npixel), orthographic similarity (OLD20), and word frequency (WF), for four different languages. (d) Representational similarity matrices (RSM; Kriegeskorte, Mur, & Bandettini, 2008) for original word images (left panel) and orthographic prediction error images (central panel). Each similarity matrix reflects the correlations among the gray values of all 3,110 words (in total 9,672,100 correlations per matrix), with words sorted alphabetically (color scale equivalent to the one used in panel b). The right panel shows the correlation between word- and orthographic prediction error-based RSMs. Each dot represents a position on the similarity matrix, allowing us to assess the relationship between the similarity values derived from the physical input and from the prediction error image. The high correlation shown here indicates that the similarity structure, or in other words, the discriminability present in the physical input, is retained in the prediction error images.

#### Accounting for Word Recognition Behavior

German lexical decision task 1: 800 five-letter nouns and 800 five-letter nonwords (400 pronounceable pseudowords, 400 unpronounceable non-words/consonant strings) were presented in pseudorandomized order (Experiment Builder software, SR-Research, Ontario, Canada; black on white background; Courier-New font; .3° of visual angle per letter; 21″ LCD monitor with 1,024 × 768 resolution and 60Hz refresh rate), preceded by ten practice trials. Participants judged for each letter string, whether it was a word or not using a regular PC keyboard, with left and right arrow keys for words and non-words, respectively. Before stimulus presentation, two black vertical bars (one above and one below the vertical position of the letter string) were presented for 500 ms, and letter strings were displayed until a button was pressed. Response times were measured in relation to the stimulus onset. German lexical decision task 2 is a replication of this experiment, but also presenting words and non-words, including visual noise. In three blocks, items with 0%, 20%, or 40% added visual noise were presented (140 items per block; 70 five-letter words and 70 nonwords, of which 36 were pseudowords and 34 were consonant clusters). Visual noise was added by replacing the respective number of pixels (for details see Gagl, Hawelka, Richlan, Schuster, & Hutzler, 2014).

Linear mixed model (LMM) analysis implemented in the lme4 package (Bates, Mächler, Bolker, & Walker, 2015) of the R statistics software was used for analyzing lexical decision data, as LMMs are optimized for estimating statistical models with crossed random effects for items and participants. These analyses result in effect size estimates with confidence intervals (SE) and a *t*-value. Following standard procedures, *t*-values larger than 2 are considered significant since this indicates that the effect size ±2 SE does not include zero (Kliegl, Wei, Dambacher, Yan, & Zhou, 2011). For the presentation in Fig. 3a,b,d,e,g,h,k,l, co-varying effects were removed by the *keepef* function of the *remef* package (Hohenstein & Kliegl, 2014/2017). We considered all response times below 300 ms as too fast given the minimal physiological constants (i.e., 70 ms for information arriving in V1), and all response times above 4.000 ms as too slow, likely due to machine error (e.g., too soft keypress); these trials were removed from further analyses. After that, response times were log-transformed, which accounts for the ex-Gaussian distribution of response times. The orthographic prediction error and the number of pixels parameters were centered and normalized by R’s *scale*() function in order to optimize LMM analysis. Linear effect models with crossed random effects of participants and items are fitted based on on single-trial data (i.e., without aggregating prior to statistical analysis), so those error responses (i.e., by explicitly modeling the effect of errors as fixed effect) and outliers are accounted for in the model and do not have to be excluded prior to the analysis (for details see Baayen, Davidson, & Bates, 2008).

**Figure 3.**
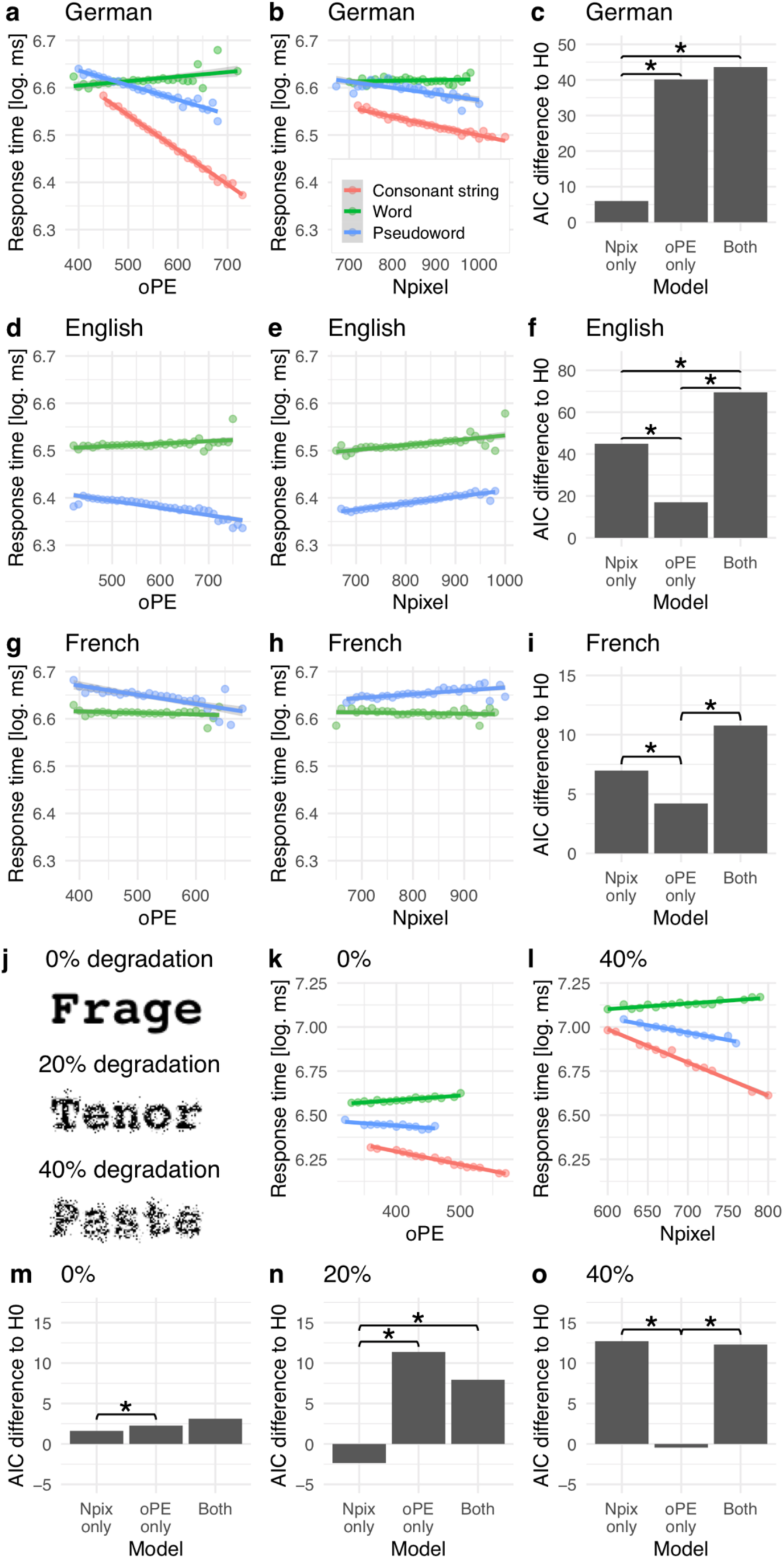
Lexical (i.e., Word/non-word) decision task behavior. (a) Orthographic prediction error (oPE) and (b) number of pixels (Npixel) effects on response times in a lexical decision task (German nouns, five letters length; overall error rate 7.4%; see Supplemental Table 1 for detailed statistical analysis). Green lines show the effects for words, blue lines for pseudowords (pronounceable non-words), and red lines for consonant strings (unpronounceable non-words). Dots represent mean reaction time estimates across all participants, separated into bins of oPE (width of 10) and stimulus category, after excluding confounding effects. (c) Results from model comparisons. First, a null model was established with only word/non-word status and word frequency as predictors. Subsequently, a model adding only the oPE predictor, a model adding only the Npixel predictor, and one model adding both predictors to the null model, were compared to the null model. Note that also the interaction terms with the word/non-word parameter were included. The Akaike Information Criterion (AIC) for the difference to the null model is shown for each model. A positive value represents an increase in model fit; asterisks mark significant differences (p < .05 Bonferroni corrected for multiple comparisons; 6 comparisons, three in relation to the null model and three, marked with asterisks, comparing the alternative models; corrected significance threshold p < .0083). (d-f) Analogous results for English and (g-i) for French lexical decision tasks. Visual noise experiment: (j) Example stimuli representing the three visual noise levels. (k) Orthographic prediction error effect (oPE) when no noise was applied, replicating the first study presented in *a* (error rate: 6%). (l) Number of pixels effect (Npixel) in the condition where the noise was strongest (error rate: 33%). (m-o) Model comparisons, including the full models and the models with oPE and Npixels only for each of the noise levels. Note that for the noise study, AIC comparisons were Bonferroni corrected for nine comparisons (corrected significance threshold p < .0055).

To assess model fit, we used the Akaike Information Criterion (AIC), as it penalizes model complexity (Akaike, 1973). Note that we have explicitly decided against reporting the percentage of variance explained (which is a commonly used metric for evaluating the contribution of specific parameters to regression models), as this metric does not take into account model complexity. Considering model complexity explicitly is important, as each additional parameter in a model per se increases the amount of explained variance (up to the extreme case of complete overfitting), but reduces the interpretability of the model (see Burnham & Anderson, 2004 for an in-depth discussion). A fair model evaluation, therefore, requires to relate explained variance to model complexity, which the AIC assures.

#### Cortical Representation of the Orthographic Prediction Error

Sixty five-letter words and 180 pseudowords were presented in a pseudorandom order (yellow Courier New font on gray background; 800 ms per stimulus; ISI 2,150 ms) as well as 30 catch trials consisting of the German word *Taste* (button), instructing participants to press the response button. Catch trials were excluded from the analyses. All items consisted of two syllables and were matched on OLD20 (Yarkoni et al., 2008) and mean bigram frequency between conditions. To facilitate estimation of the hemodynamic response, an asynchrony between the TR (2,250 ms) and stimulus presentation (onset asynchrony: 2,150 + 800 ms) was established and 60 null events were interspersed among trials; a fixation cross was shown during inter-stimulus intervals and null events. The sequence of the presentation was determined by a genetic algorithm (Wager & Nichols, 2003), which optimized for maximal statistical power and psychological validity. The fMRI session was divided into two runs with a duration of approximately 8 min each.

A Siemens Magnetom TRIO 3-Tesla scanner (Siemens AG, Erlangen, Germany) equipped with a 32-channel head-coil was used for functional and anatomical image acquisition. The BOLD signal was acquired with a T_2_*-weighted gradient-echo echo-planar imaging sequence (TR = 2,250 ms; TE = 30 ms; Flip angle = 70°; 86 x 86 matrix; FoV = 192 mm). Thirty-six axial slices with a slice thickness of 3 mm and a slice gap of 0.3 mm were acquired in descending order within each TR. In addition, for each participant a gradient echo field map (TR = 488 ms; TE 1 = 4.49 ms; TE 2 = 6.95 ms) and a high-resolution structural scan (T_1_-weighted MPRAGE sequence; 1 x 1 x 1.2 mm) were acquired. Stimuli were presented using an MR-compatible LCD screen (NordicNeuroLab, Bergen, Norway) with a refresh rate of 60 Hz and a resolution of 1,024×768 pixels.

SPM8 software (http://www.fil.ion.ucl.ac.uk/spm), running on Matlab 7.6 (Mathworks, Inc., MA, USA), was used for preprocessing and statistical analysis. Functional images were realigned, unwarped, corrected for geometric distortions by use of the FieldMap toolbox, and slice-time corrected. The high-resolution structural image was pre-processed and normalized using the VBM8 toolbox (http://dbm.neuro.uni-jena.de/vbm8). The image was segmented into gray matter, white matter, and CSF compartments, denoised, and warped into MNI space by registering it to the DARTEL template of the VBM8 toolbox using the high-dimensional DARTEL registration algorithm (Ashburner, 2007). Functional images were co-registered to the high-resolution structural image, which was normalized to the MNI T_1_ template image, and resulting normalization parameters were applied to the functional data, which were then resampled to a resolution of 2×2×2 mm and smoothed with a 6 mm FWHM Gaussian kernel.

For statistical analysis, we first modeled stimulus onsets with a canonical hemodynamic response function and its temporal derivative, including movement parameters from the realignment step and catch trials as covariates of no interest, a high-pass filter with a cut off of 128 s, and an AR(1) model (Friston et al., 2002) to correct for autocorrelation. The fMRI analysis involved a predictor with item onsets as events of interest; besides, we introduced a continuous, item-specific predictor reflecting variation on the orthographic measure of interest. Please note that we ran separate analyses for the oPE simulations, OLD20, lexicality, and the oPE by lexicality interaction. The main reason for this procedure is that regression models for the fMRI analysis, are less flexible than the crossed-random effect design we applied for the analysis of behavioral and EEG data. First, i.e., first-level analysis, the BOLD response for the oPE predictor is modeled across multiple trials for each participant. Second, i.e., on the second level, statistical tests, for each voxel, are implemented across participants. This procedure limits the variance to the inter-subject component as the inter-stimulus component is removed on the first level of the analysis. Therefore, one is limited with the introduction of multiple, correlated predictors (e.g., the oPE and OLD predictors; see Fig. 2b,c), as the amount of variance to be explained is already limited, to come up with a reasonable result. One benefit of fMRI is the high number of regions, i.e., voxels, we can estimate our analysis. Meaning, it is expected that two processes that are described by two correlated predictors should be implemented in similar regions and vice versa. Thus, we opted for a single predictor analysis allowing that predictor to explain as much variance as possible. In case regions overlap, we compare peak voxel T-values to evaluate which (i.e., the one with the higher effect size) predictor is adequate. Group level effects are implemented as one-sided t-tests with a voxel-level threshold of *p* < .001 uncorrected and a cluster-level correction for multiple comparisons (*p* < .05 family-wise error corrected). Where peak effects survived voxel-level family-wise error correction, this is additionally reported. fMRI results are visualized using ‘glass brain’ figures from the nilearn python package (Abraham et al., 2014), and anatomical labels for activation clusters were extracted with the AtlasReader python package (Notter et al., 2019).

#### Timing of the Orthographic Prediction Error

200 five-letter words, 100 pseudowords, and 100 consonant strings (nonwords) were presented for 800 ms (black on white background; Courier-New font, .3° of visual angle per letter), followed by an 800 ms blank screen and a 1,500 ms hash mark presentation, which marked an interval in which the participants were instructed to blink if necessary. In addition, 60 catch trials (procedure as described for the fMRI study) were included in the experiment. Stimuli were presented on a 19″ CRT monitor (resolution 1,024 × 768 pixels, refresh rate 150Hz), and were preceded by two black vertical bars presented for 500 - 1,000 ms to reduce stimulus onset expectancies.

EEG was recorded from 64 active Ag/Ag-Cl electrodes (extended 10-20 system) using an actiCAP system (BrainProducts, Germany). FCz served as common reference and the EOG was recorded from the outer canthus of each eye as well as from below the left eye. A 64-channel Brainamp (BrainProducts, Germany) amplifier with a 0.1–1,000 Hz bandpass filter sampled the amplified signal with 500Hz. Electrode impedances were kept below 5kΩ. Offline, the EEG data were re-referenced to the average of all channels. EEG data were preprocessed using MNE-Python (Gramfort et al., 2014), including high (.1 Hz) and low pass (30 Hz) filtering and removal of ocular artifacts using ICA (Delorme, Sejnowski, & Makeig, 2007). For each subject, epochs from 0.5 s before to 0.8 s after word onset were extracted and baselined by subtracting the pre-stimulus mean, after rejecting trials with extreme (>50 μV peak-to-peak variation) values. Multiple regression analysis, with the exact same parameters as for the behavioral evaluation (orthographic prediction error, number of pixels, word/non-word, and the interactions with the word/non-word distinction), was conducted and a cluster-based permutation test (Maris & Oostenveld, 2007) was used for significance testing. 1,024 label permutations were conducted to estimate the distribution of thresholded clusters of spatially and temporally (i.e., across electrodes and time) adjacent time points under the null hypothesis. All clusters with a probability of less than an assumed alpha value of .05 under this simulated null hypothesis were considered statistically significant. For the presentation of effect patterns (line and box-plots) in Fig. 6, co-varying effects were removed by the *keepef* function of the *remef* package (Hohenstein & Kliegl, 2014/2017).

#### Application to handwriting

To provide a first demonstration that prediction-based reading can also be generalized to less standardized situations, we explored whether the oPE of different persons’ handwriting is associated with their readability. We obtained handwriting samples (26 upper and 26 lower case letters; 10 common German compound words, 10-24 letters long) from 10 different writers (see Fig. 6a,b, for examples). The single letters were scanned and centered within a 50×50 pixel image. These images were used to estimate, for each writer’s data separately, pixel-by-pixel predictions for upper and lower-case letters (see also Fig. 6a,b), analogous to the procedures described above and in Fig. 1b. Subsequently, these predictions were subtracted from each letter of the alphabet within the respective writer’s samples (matrix subtraction; analogous to Formula 1). We chose this procedure (as opposed to the word-based calculation of oPEs for the remaining parts of the study) since the alignment of letters is much easier compared to words. In contrast to computer fonts, the correlation of the orthographic prediction error and the particular item’s number of pixels was high (r = .98). We normalized the orthographic prediction error by a division with the respective pixel count to compensate for this confound. Based on the mean of the corrected prediction errors per letter, we obtained two oPE estimates, one for upper and one for lower case letters, for each handwriting. Readability ratings (5-point Likert scale) were obtained from 38 participants (27 females; mean age 25 years) by presenting all ten versions of all ten handwritten words, in addition to the identical word in the computerized script. We used an LMM analysis to estimate the effect of the orthographic prediction error (i.e., only fixed effect) on the readability ratings. Also, we include the random effect of participants on the intercept of the orthographic prediction error slope. Besides, to investigate the relationship of input signal of handwritten letters directly to the predictions and prediction errors derived from the input signal, we implemented a LMM analysis that correlated the orthographic prediction error (Fig. 6c-d) with the mean prediction strength (i.e., mean of the values extracted from the prediction matrix), number of all non-white pixels (both scaled), and letter case. Besides, we added the random effect on the intercept for each handwriting to the model.

## Results

### Lexicon-based Characterization of the Orthographic Prediction Error

Cognitive psychologists have developed several quantitative measures to characterize words (Brysbaert et al., 2011; Coltheart et al., 1977; Yarkoni et al., 2008), mostly derived from large text corpora and psycholinguistic word databases (like the SUBTLEX database; (Heuven et al., 2014; Keuleers, Brysbaert, et al., 2010). Fig. 2a describes some of the most widely-used lexical word characteristics that are relevant for the present work and gives examples to illustrate them. Abundant empirical research demonstrates that these lexicon-based word characteristics are predictive of different aspects of reading behavior (e.g., Balota, Cortese, Sergent-Marshall, Spieler, & Yap, 2004; Rayner, 2009). Accordingly, understanding how the orthographic prediction error, derived from the implemented PEMoR (see Fig. 1), relates to these measures can provide an important first indication if and how this optimized and supposedly pre-lexical perceptual signal is involved in word recognition.

A hierarchical cluster analysis (Wei & Simko, 2017) indicates that across all words, the orthographic prediction error (i.e., the sum of all gray values after subtracting the knowledge-based prior from the actual stimulus image; cf. Fig. 1d and Methods) clusters with several measures that are commonly interpreted as reflecting orthographic word properties (Fig. 2b). More specifically, the oPE clusters with widely-used (psycho-) linguistic characteristics reflecting the (non-) uniqueness of words in terms of their orthographic similarity to other words (i.e., the number of Coltheart neighbors; Coltheart et al., 1977) or their orthographic distance (OLD20; Yarkoni et al., 2008; cf. Fig. 2a) and letter co-occurrence statistics (i.e., bi-, tri- and quadrigram frequencies; cf. Fig. 2a). Note that these measures describe the statistics of letters and letter combinations in all words retrieved from a lexicon database (Keuleers, Brysbaert, et al., 2010). In cognitive psychological research, these measures are consistently associated with the first, i.e., orthographic, stages of processing written words before lexical access (Coltheart et al., 2001; Grainger & Jacobs, 1996). The significant correlations between the oPE and these lexical-orthographic measures are particularly interesting as they demonstrate that a neurophysiologically inspired transformation of the visual stimulus, i.e., the here-proposed orthographic prediction error (oPE), is meaningfully related to orthographic properties of words as derived from lexicon-based statistics. Crucially, this is achieved while (a) reducing the to-be-processed signal by more than 30% and (b) at the same time retaining the ability of discriminating the word identities, as indicated by a strong correlation of *r* = .87 between the representational similarity matrices (Edelman, 1998; Kriegeskorte et al., 2008) of the word and orthographic prediction error images (Fig. 2d). This latter result indicates that the representational similarity structure, or in other words, the discriminability between items, is preserved after deriving the oPE from the sensory input as proposed by the PEMoR.

In contrast, the orthographic prediction error is not correlated with the frequency of occurrence of a word in a language (Fig. 2b). The word frequency effect (i.e., faster response times to more frequently occurring words) typically indicates the difficulty of accessing word meaning based on an already-decoded orthographic access code (Coltheart et al., 2001). This dissociation between the orthographic prediction error and word frequency replicates across languages (Fig. 2c) and is much more pronounced for the orthographic prediction error than for the so-far dominant measures of orthographic similarity (OLD20) and orthographic neighborhood (Fig. 2b). Only trigram and quadrigram frequency were significantly correlated with the raw pixel count of words (which represents an approximation of the full physical signal contained in the stimuli (Fig. 2b). This dissociation of the correlation structure of the prediction error and the pixel count provides the first evidence that the neurophysiologically inspired orthographic prediction error is more important for a mechanistic understanding of reading than the full physical input contained in a printed word as assumed in the baseline model representing the full pixel count of the input image.

#### Accounting for Word Recognition Behavior

As a next empirical test of the prediction error model of reading (PEMoR), we evaluated how well the orthographic prediction error performs in accounting for behavior in an established and widely-used word recognition task, i.e., the lexical decision task. Thirty-five human participants were asked to decide as fast as possible by button press whether written letter-strings (presented on the computer screen; 1,600 items; 5 letters length; language: German) were words or not. Remember that the orthographic prediction error represents the deviance of a given letter-string from our knowledge-based orthographic expectation, and thus how unlikely it is that the given letter-string is a word. Accordingly, participants should be fast in identifying letter-strings with low orthographic prediction error as words but slow in identifying words with a high orthographic prediction error, and fast in rejecting non-words with a high orthographic prediction error.

Fig. 3a shows exactly this pattern of response times, i.e., a word/non-word by orthographic prediction error interaction (linear mixed model/LMM estimate: 0.03; SE = 0.01; t = 5.0; see Methods for details on linear mixed effects modeling and Supplemental Table 1 for detailed results). No significant interaction or fixed effect of the number of pixels estimate (i.e., the sum of all pixels contained in a word) was found (Fig. 3b; Interaction: estimate: 0.00; SE = 0.01; t = 0.0; Fixed effect: −0.01; SE = 0.00; t =1.8). To directly compare if the response times are more adequately described by the PEMoR or the baseline model representing the full pixel count of the input image, we performed an explicit model comparison (see Methods for details) of four models, i.e.,(i) a full model, including as predictors the orthographic prediction error and the number of pixels, (ii) a pure prediction error model, (iii) a pure number of pixels model, and (iv) a null model without any of the two predictors. Fig. 3c shows that, in contrast to the null model, the three alternative models showed higher model fits (all χ^2^’s > 9.9; all *p*’s < .007; Bonferroni corrected *p* threshold: 0.0083), but this increase was significantly larger for the models including the orthographic prediction error. In addition, the model including only the orthographic prediction error explained substantially higher amounts of variance when compared to the model including only the number of pixels parameter (AIC difference: 34; χ^2^(0) = 34.2; p < .001) with no substantial further increase for the combined model (AIC difference: 3; χ^2^(2) = 7.2; p = .02). This finding indicates that for visual word recognition in German, the baseline model representing the full pixel count of the input image explains substantially less variance in word recognition behavior than the PEMoR.

Additionally including orthographic distance (OLD20; Yarkoni et al., 2008) as predictor improved the model fit further (AIC difference comparing the full model with and without OLD20: 104; χ^2^(2) = 105.8; p < .001) but did not affect the significance of the word/non-word-by-orthographic prediction error interaction (estimate of the interaction effect after including additional parameters: 0.03; SE = 0.01; *t* = 5.2). This finding indicates that despite its correlation with other orthographic measures (Fig. 2b, c), the orthographic prediction error accounts for unique variance components in word recognition behavior that cannot be explained by other word characteristics.

We also replicate this interaction when calculating the orthographic prediction error using a length-unspecific visual-orthographic prediction (i.e., based on all ∼190,000 German words from the SUBTLEX database; Brysbaert et al., 2011; 2 - 36 letters length; cf. Fig. 1e; LMM estimate of interaction effect: 0.03; SE = 0.01; *t* = 4.5; for replication in English and a more extensive investigation of the interaction effect for multiple word lengths see Supplemental figure 1a). Interestingly, length-specific and length-unspecific orthographic prediction errors are highly correlated (e.g., German: *r* = .97), suggesting strongly that the prediction-based word recognition process proposed by the PEMoR model is independent of word length constraints. This finding is in line with evidence from natural reading, which shows that one can extract low-level visual features like the number of letters from the parafoveal vision before fixating the word (Cutter, Drieghe, & Liversedge, 2014; Gagl et al., 2014; Schotter, Angele, & Rayner, 2012). The use of a fixed word length in our German lexical decision experiment is therefore not necessarily artificial since in natural reading, word length is perceived before fixation. In sum, these results demonstrate that the orthographic prediction error is meaningfully related to word recognition behavior and independent of word length. Finally, we also investigated whether the morphology of the words may contribute to lexical decision performance. To this end, we included the number of morphemes (mean: 1.3; range 1-2) as a further predictor into best-fitting model (see ii above). The number of morphemes was only weakly correlated with the oPE (r = .16), had an effect on decision times (Estimate: −0.03; SE = 0.01; t = 3.5), but did not affect the significance of the word/non-word-by-orthographic prediction error interaction (estimate of the interaction effect after including additional parameters: 0.03; SE = 0.01; *t* = 4.9).

##### Generalization across languages

The interaction effect between lexicality (word/non-word status) and orthographic prediction error could be replicated in two open datasets from other languages, i.e., British English (Keuleers et al., 2012; 78 participants and 8,488 five-letter words/non-words: Fig. 3d; estimate: 0.008; SE = 0.002; *t* = 4.2) and French (Ferrand et al., 2010; 974 participants and 5,368 five-letter words/non-words: Fig. 3g; estimate: 0.005; SE = 0.002; *t* = 2.0); see also Supplemental Figure 2 for two further datasets from Dutch and Supplemental Table 1 for detailed results. However, in contrast to German, in both datasets we also found a significant effect of the number of pixels parameter (Fig. 3e,h; British: fixed effect: 0.008; SE = 0.001; *t* = 6.7; French: interaction with word/non-word status: −0.007; SE = 0.002; *t* = 3.0). In terms of model comparison, the pattern derived from German, i.e., the greatest increase in model fit when including the orthographic prediction error, could not be recovered for English and French. Rather, we found that the role of the number of pixels parameter for describing the response times was larger than in German (see Fig. 3f,i). Still, the combined model showed the best model fit in all three languages (Fig. 3f,i; oPE only vs. full model: AIC difference English: 52; χ^2^(2) = 56.5; p < .001; French: 6; χ^2^(2) = 10.5; p = .005; Npixel only vs. full model: AIC difference English: 24; χ^2^(2) = 28.6; p < .001; French: 3; χ^2^(2) = 7.8; p = .02; Bonferroni corrected *p* threshold: 0.0083) indicating that both the orthographic prediction error and the number of pixels parameter are relevant in explaining lexical decision behavior. To summarize, for English and French, model comparisons showed that in addition to the prediction error, the parameter reflecting more directly the physical stimulus input explained a greater amount of variance than in German. Nevertheless, in all three languages, the orthographic prediction error explained unique variance components, which further supports its relevance for understanding visual word recognition. Future research should aim at clarifying the differential reliance on the bottom-up input itself in different languages (but see also the next section for a potential explanation).

##### Word recognition behavior under conditions of visual noise

We speculated that the more significant role for bottom-up input in the British and French datasets might result from the presence of several sources of additional perceptual variability. For example, word length changed from trial to trial (English, 2-13 letters; French, 2-19 letters) and a proportional font (Times new roman) was used in the English dataset, while we had used only five-letter words presented in a monospaced font in the German experiment (Fig. 1). Even though such unpredictable perceptual variation, without any doubt, is not the standard case in naturalistic reading (i.e., through the integration of visual information from parafoveal vision; see above), in a single word reading paradigm it reduces the ability to predict visual features of upcoming stimuli and thus unnaturally decreases the performance of our model. For example, using a proportional-spaced font removes structure (e.g., the letter separation) both in the sensory input and the orthographic prior (e.g., see Supplemental Figure S2g), which results in less precise predictions and greater prediction errors. This reduction in prediction strength, in turn, increases the correlation between the number of pixels in the input image and the derived orthographic prediction error (cp. Monospace font: r = .05 vs. proportional font: r = .49, both in German; see Supplemental Figure S2g). In the face of this, it is particularly noteworthy that the orthographic prediction error, as proposed here, remained a highly relevant predictor in the English and French data set. It should also be stressed again that in natural reading, low-level visual features like word length or letter position can be picked up in parafoveal vision, so that the visual system may be able to dynamically adapt its predictions to the upcoming word (Schotter et al., 2012). Future work will, therefore, have to specify in more detail the nature of orthographic priors in naturalistic reading.

To directly test if visual word recognition relies more firmly on the bottom-up input when visual word presentation includes unpredictable perceptual variations, we conducted a second lexical decision experiment. We presented visual word stimuli with an explicit manipulation of visual noise (0% vs. 20% vs. 40% noise level) to reduce the predictability of visual features (for details see Methods section). We here applied a noise manipulation (rather than, e.g., a comparison of different fonts) since noise levels can be easily manipulated and quantified (i.e., in terms of the number of displaced pixels). In contrast, a direct comparison of fonts is more difficult because the contrast of proportional vs. mono-spaced font is confounded with many other visual differences like total stimulus width (Hautala, Hyönä, & Aro, 2011; Marinus et al., 2016). In addition, the 0% noise stimuli allowed us to replicate our original behavioral finding. Figure 3j shows examples of the word stimuli under different noise levels.

We found, in general, that response times and errors increased with the amount of noise that was applied to the visual-orthographic stimuli (0%: response time/RT: 613 ms, 6% errors; 20%: RT: 739 ms, 12% errors; 40%: RT: 1,105 ms, 33% errors; compare also Fig. 3k and l). When no noise was applied we replicated our first study (cp. Fig. 3k and a) with a significant interaction between the orthographic prediction error and the word/non-word factor (estimate: 0.05; SE = 0.02; *t* = 2.3; see Supplemental Table 1 for detailed results). As in the first experiment, no effect or interaction was found for the number of pixels parameter. With 20% noise, we still could identify a fixed negative effect of the orthographic prediction error (estimate: −0.06; SE = 0.02; *t* = 3.3) however without a significant interaction pattern. Also, the fixed effect of the number of pixels was not significant. With 40% noise, however, no significant effect of the orthographic prediction error could be found, but as expected from the above discussion of noise effects, we observed now a significant fixed effect of the number of pixels parameter, as well as an interaction with word/non-word status (Fig. 3l; estimate: 0.08; SE = 0.03; *t* = 2.9). A similar impression can be obtained from the model fit results showing that including the orthographic prediction error resulted in significantly higher model fits for 0% and 20% noise conditions compared to models in which only the number of pixels predictor was included (see Fig. 3m,n; 0% AIC difference: 1; χ^2^(0) = 1; p < .001; 20% AIC difference: 13; χ^2^(0) = 13.7; p < .001). With 40% noise, inclusion of the number of pixels parameter resulted in a higher model fit (see Fig. 3o; AIC difference: 13; χ^2^(0) = 13.2; p < .001), but including the orthographic prediction error had essentially no effect.

In sum, the behavioral experiments reported in this section demonstrate that the orthographic prediction error contributes substantially to visual word recognition. We find that the PEMoR is highly relevant when the visual information presented in the lexical decision tasks is restricted in variability and, therefore, more predictable (i.e., all words with the same number of letters). Better predictability of visual features results in greater reliance on the orthographic prediction error compared to the pure bottom-up sensory input. In contrast, when the perceptual variability (i.e., complexity) increases, e.g., due to a variation of the number of letters, proportional fonts (e.g., as in the case of the English study), or visual noise, the full (i.e., non-optimized) bottom-up signal becomes more important for explaining visual word recognition behavior. Thus, the behavioral evidence indicates that efficient neuronal coding is used when the perceptual properties of the letter strings can be predicted. In case the perceptual properties are highly variable, predictive processing is hampered as only weak predictions can be formed, suggesting that the baseline model representing the full bottom-up signal is a “fallback” strategy to as an approximation in effortful reading conditions.

#### Cortical Representation of the Orthographic Prediction Error

The PEMoR assumes that the orthographic prediction error is estimated at an early stage of word recognition, i.e., in the visual-perceptual system and before word meaning is accessed and higher-level linguistic representations of the word are activated. We accordingly hypothesized that brain systems involved in computing or representing the orthographic prediction error should be driven by this optimized representation of the sensory input independent of the item’s word/non-word status (i.e., for words and non-words alike). Localizing the neural signature of the orthographic prediction error in the brain during word/non-word recognition, thus, is a further critical test of the PEMoR. Of note, a strict bottom-up model of word recognition (and perception in general) would make a different prediction, i.e., that activation in visual-sensory brain regions should be driven by the full amount of physical signal in the percept (Goodyear & Menon, 1998; Henrie & Shapley, 2005). Processes that take place after word-identification, i.e., that involve higher levels of linguistic elaboration can only operate on mental representations of words, so that brain regions involved in these later stages of word processing should distinguish between words and non-words.

We tested these hypotheses about the localization of the orthographic prediction error by measuring BOLD activation changes using functional MRI, while 39 participants silently read words (German nouns) and pronounceable non-words (i.e., pseudowords), in randomized order (see Methods for details). We identified three left- and two right-hemispheric brain regions in the occipital cortex that showed higher levels of activation when reading items with higher orthographic prediction error – both for words and non-words (Fig. 4a and Table 1). Prior research (Dehaene & Cohen, 2011; Dehaene et al., 2005) has identified a region in the mid-portion of the left occipito-temporal cortex as critical for reading: the visual word form area. Consistent with our hypothesis, all five activation clusters representing the orthographic prediction error are located posterior to this so-called visual word form area (Dehaene & Cohen, 2011), which supports our claim of an ‘early’ perceptual role for the orthographic prediction error signal before word identification. More recently, (Lerma-Usabiaga et al., 2018) proposed a distinction of perceptual (i.e., all y-coordinates < - 60) and lexical processing regions of the left occipito-temporal cortex. All oPE-related activation clusters were located posteriorly within the perceptual processing regions (see Table 1). To compare the BOLD results for the oPE with a more established measure of orthographic similarity, we conducted the identical analysis with OLD20, rather than the oPE, as a parametric predictor. In this control analysis, we found only one left-hemispheric cluster within these perceptual processing regions, and this cluster showed a lower peak activation (i.e., lower t value at the cluster’s peak voxel) compared to the oPE clusters. Importantly, no brain areas showed activity dependent on the pure amount of bottom-up signal in the percept (i.e., the number of pixels parameter).

**Figure 4.**
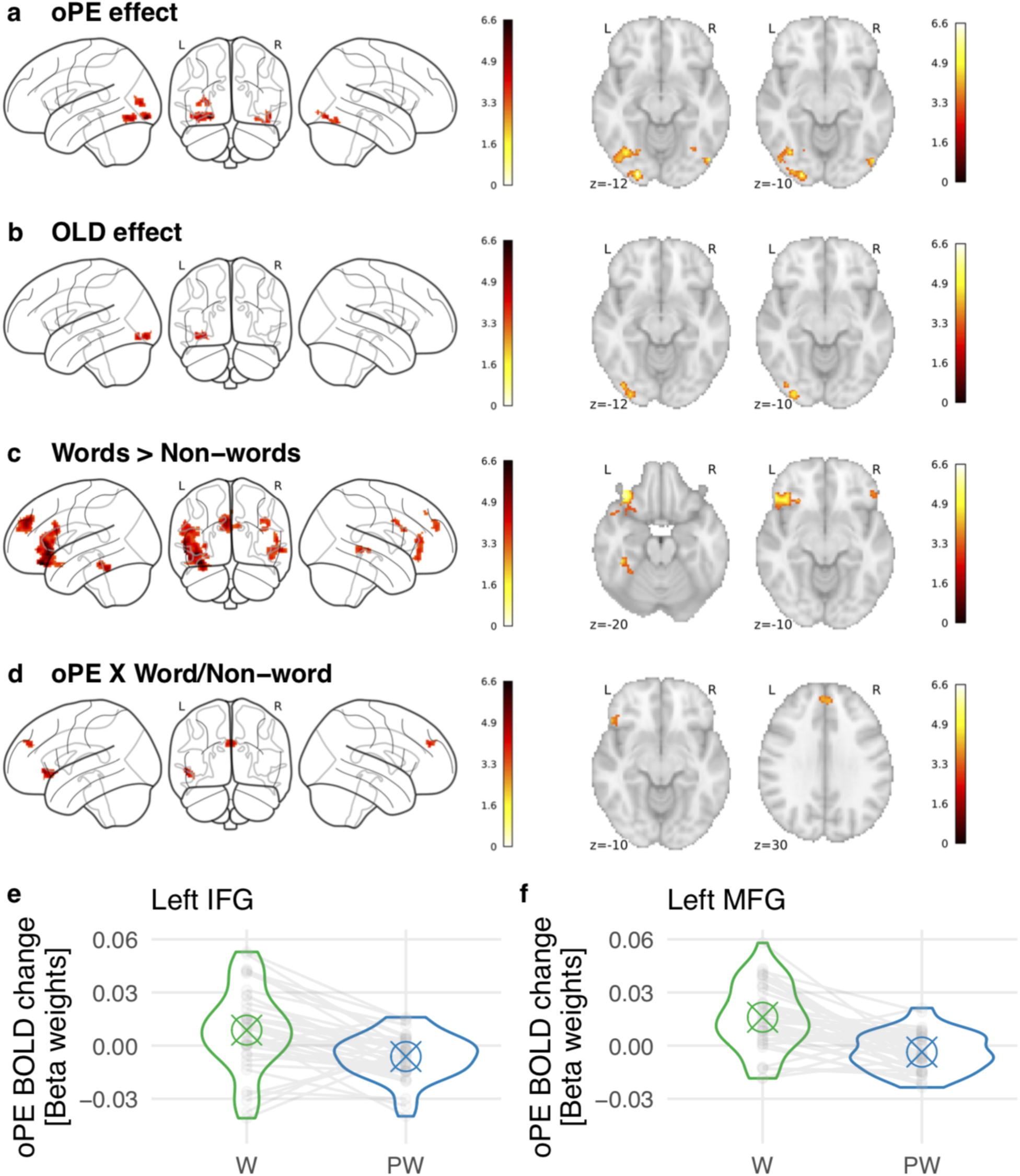
fMRI results demonstrating the neuroanatomical localization of orthographic prediction error effects. BOLD activation during silent reading (see Materials and Methods for further details, and Table 1 for exact locations of activation effects). (a) An analysis demonstrating a positive orthographic prediction error (oPE) effect in bilateral occipital activation-clusters. This regression analysis used item-specific oPE values as a covariate, independent of stimulus condition, and shows brain regions with greater activity for letter strings characterized by a higher oPE, independent of stimulus type. (b) The same analysis for the OLD20 parameter (Yarkoni et al., 2008), which represents an established measure of orthographic similarity. (c) Clusters of higher BOLD activation for words than for non-words. (d) Two frontal activation clusters showing an oPE by word/non-word interaction, i.e., positive and negative beta weights representing the oPE effect for each subject, separated for words (W) and non-words (in this case only pseudowords; PW), respectively. Points within the violin figures represent the individual subjects’ beta weights and the crossed circle symbol represents the mean beta weights (crossed circle) for the peak of (e) the left IFG cluster, i.e., x = −52, y = 32, z = −4, and (f) the left MFG cluster, i.e., x = −4, y = 48, z = 28; lines connect word and non-word betas from each individual. These values demonstrate a positive oPE effect on BOLD for words and a negative oPE effect on BOLD for non-words. No effects of the number of pixels per word were found. Threshold voxel-level: p<.001 uncorrected; cluster level: p<.05 family-wise error corrected.

**Table 1.**
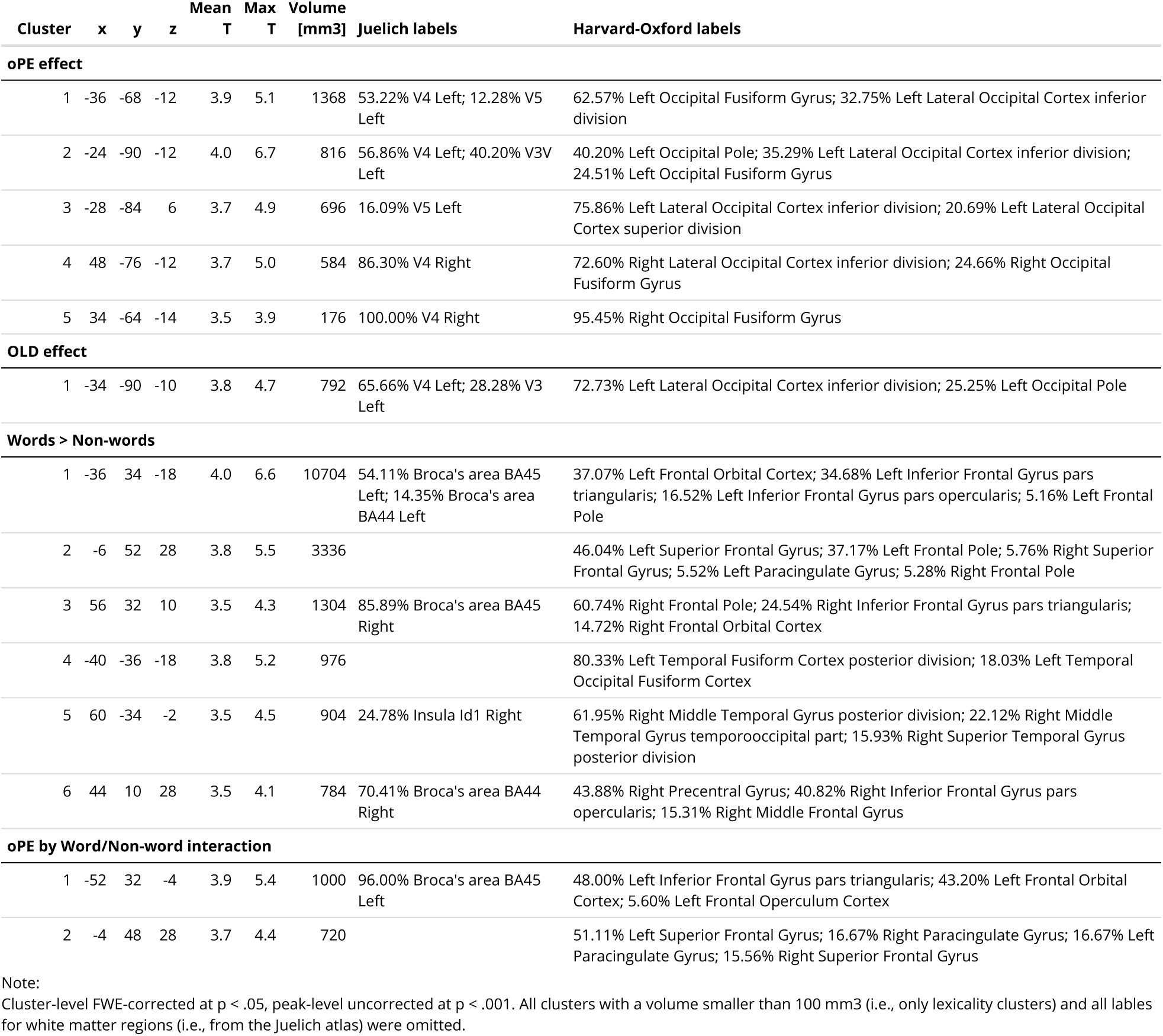
Reliable activation clusters from the fMRI evaluation with respective anatomical labels (most likely regions from the Juelich and Harvard-Oxford atlases including % overlap), cluster size (in mm3), and peak voxel coordinates (MNI space).

| Cluster | x | y | z | Mean T | Max T | Volume [mm<sup>3</sup>] | Juelich labels | Harvard-Oxford labels |
| --- | --- | --- | --- | --- | --- | --- | --- | --- |
| **oPE effect** |  |  |  |  |  |  |  |  |
| 1 | -36 | -68 | -12 | 3.9 | 5.1 | 1368 | 53.22% V4 Left; 12.28% V5 Left | 62.57% Left Occipital Fusiform Gyrus; 32.75% Left Lateral Occipital Cortex inferior division |
| 2 | -24 | -90 | -12 | 4.0 | 6.7 | 816 | 56.86% V4 Left; 40.20% V3V Left | 40.20% Left Occipital Pole; 35.29% Left Lateral Occipital Cortex inferior division; 24.51% Left Occipital Fusiform Gyrus |
| 3 | -28 | -84 | 6 | 3.7 | 4.9 | 696 | 16.09% V5 Left | 75.86% Left Lateral Occipital Cortex inferior division; 20.69% Left Lateral Occipital Cortex superior division |
| 4 | 48 | -76 | -12 | 3.7 | 5.0 | 584 | 86.30% V4 Right | 72.60% Right Lateral Occipital Cortex inferior division; 24.66% Right Occipital Fusiform Gyrus |
| 5 | 34 | -64 | -14 | 3.5 | 3.9 | 176 | 100.00% V4 Right | 95.45% Right Occipital Fusiform Gyrus |
| **OLD effect** |  |  |  |  |  |  |  |  |
| 1 | -34 | -90 | -10 | 3.8 | 4.7 | 792 | 65.66% V4 Left; 28.28% V3 Left | 72.73% Left Lateral Occipital Cortex inferior division; 25.25% Left Occipital Pole |
| **Words > Non-words** |  |  |  |  |  |  |  |  |
| 1 | -36 | 34 | -18 | 4.0 | 6.6 | 10704 | 54.11% Broca's area BA45 Left; 14.35% Broca's area BA44 Left | 37.07% Left Frontal Orbital Cortex; 34.68% Left Inferior Frontal Gyrus pars triangularis; 16.52% Left Inferior Frontal Gyrus pars opercularis; 5.16% Left Frontal Pole |
| 2 | -6 | 52 | 28 | 3.8 | 5.5 | 3336 |  | 46.04% Left Superior Frontal Gyrus; 37.17% Left Frontal Pole; 5.76% Right Superior Frontal Gyrus; 5.52% Left Paracingulate Gyrus; 5.28% Right Frontal Pole |
| 3 | 56 | 32 | 10 | 3.5 | 4.3 | 1304 | 85.89% Broca's area BA45 Right | 60.74% Right Frontal Pole; 24.54% Right Inferior Frontal Gyrus pars triangularis; 14.72% Right Frontal Orbital Cortex |
| 4 | -40 | -36 | -18 | 3.8 | 5.2 | 976 |  | 80.33% Left Temporal Fusiform Cortex posterior division; 18.03% Left Temporal Occipital Fusiform Cortex |
| 5 | 60 | -34 | -2 | 3.5 | 4.5 | 904 | 24.78% Insula Id1 Right | 61.95% Right Middle Temporal Gyrus posterior division; 22.12% Right Middle Temporal Gyrus temporooccipital part; 15.93% Right Superior Temporal Gyrus posterior division |
| 6 | 44 | 10 | 28 | 3.5 | 4.1 | 784 | 70.41% Broca's area BA44 Right | 43.88% Right Precentral Gyrus; 40.82% Right Inferior Frontal Gyrus pars opercularis; 15.31% Right Middle Frontal Gyrus |
| **oPE by Word/Non-word interaction** |  |  |  |  |  |  |  |  |
| 1 | -52 | 32 | -4 | 3.9 | 5.4 | 1000 | 96.00% Broca's area BA45 Left | 48.00% Left Inferior Frontal Gyrus pars triangularis; 43.20% Left Frontal Orbital Cortex; 5.60% Left Frontal Operculum Cortex |
| 2 | -4 | 48 | 28 | 3.7 | 4.4 | 720 |  | 51.11% Left Superior Frontal Gyrus; 16.67% Right Paracingulate Gyrus; 16.67% Left Paracingulate Gyrus; 15.56% Right Superior Frontal Gyrus |

Note:
Cluster-level FWE-corrected at $p < .05$ , peak-level uncorrected at $p < .001$ . All clusters with a volume smaller than 100 mm<sup>3</sup> (i.e., only lexicality clusters) and all labels for white matter regions (i.e., from the Juelich atlas) were omitted.

Only brain regions involved in the activation of word meaning and subsequent processes should differentiate between words and non-words. We observed higher activity for words than non-words, independent of the orthographic prediction error, more anteriorly in the left temporal and prefrontal cortex (Fig. 4b and Table 1). Third, the left inferior frontal gyrus (pars triangularis) and the medial portion of the superior frontal gyrus (mSFG) mirrored the lexical decision behavior reported above, in that higher prediction errors lead to increased activation for words but decreasing activation for non-words (lexicality x oPE interaction; Fig. 4c and Table 1). The fMRI experiment, thus, supports our hypothesis that during the early perceptual stages of visual processing, i.e., presumably before accessing word meaning, an optimized perceptual signal, the orthographic prediction error, is generated and used as a basis for efficient visual-orthographic processing of written language. Only at later processing stages (in more anterior temporal and prefrontal cortices), the brain differentiates between words and non-words.

#### Timing of the Orthographic Prediction Error

While the fMRI results demonstrate a representation of the orthographic prediction error in presumably ‘early’ visual cortical regions, the temporal resolution of fMRI precludes inferences concerning the temporal sequence of cognitive processes during word recognition. Across many studies, the millisecond time resolution of EEG has helped to consistently attribute the extraction of meaning from perceived words to a time window of around 300 to 600 ms post word onset (N400 component of the event-related brain potential/ERP; Kutas & Federmeier, 2011). Visual-orthographic processes associated with the orthographic prediction error should thus temporally precede this time window, most likely to occur during the N170 component of the ERP (Barber & Kutas, 2007; Carreiras, Armstrong, Perea, & Frost, 2014; Grainger & Holcomb, 2009). To test this hypothesis, we measured EEG while 31 participants silently read words and non-words (including both pseudowords and consonant-only strings). We fitted a multiple regression model (analogous to the model used for the analysis of behavioral data) to the EEG data (Linzen & Engemann, 2017) with the orthographic prediction error, the number of pixels, word/non-word-status, and their interactions as parameters (see Materials and Methods for details).

Regression-estimated ERPs (see methods for details) show a significant effect of the orthographic prediction error on electrical brain activity between 150 and 250 ms after stimulus onset (Fig. 5a). In this early time window, letter-strings characterized by higher prediction errors elicited significantly more negative-going ERPs over posterior-occipital sensors, for both words and non-words. In line with the temporal sequence of processes inferred from their neuroanatomical localizations (i.e., fMRI results), a significant word/non-word effect then emerged between 200-570 ms (Fig. 5b), followed by an interaction between word/non-word-status and orthographic prediction error at 360-620 ms (Fig. 5c). In this interaction cluster, higher prediction errors led to more negative-going ERPs for non-words, as observed for all stimuli in the earlier time window, but showed a reverse effect for words, i.e., more positive-going ERPs for words with higher prediction errors (Fig. 5c). This pattern of opposite prediction error effects for words vs. non-words is analogous to the effects seen in lexical decision behavior and the frontal brain activation results obtained with fMRI.

**Figure 5.**
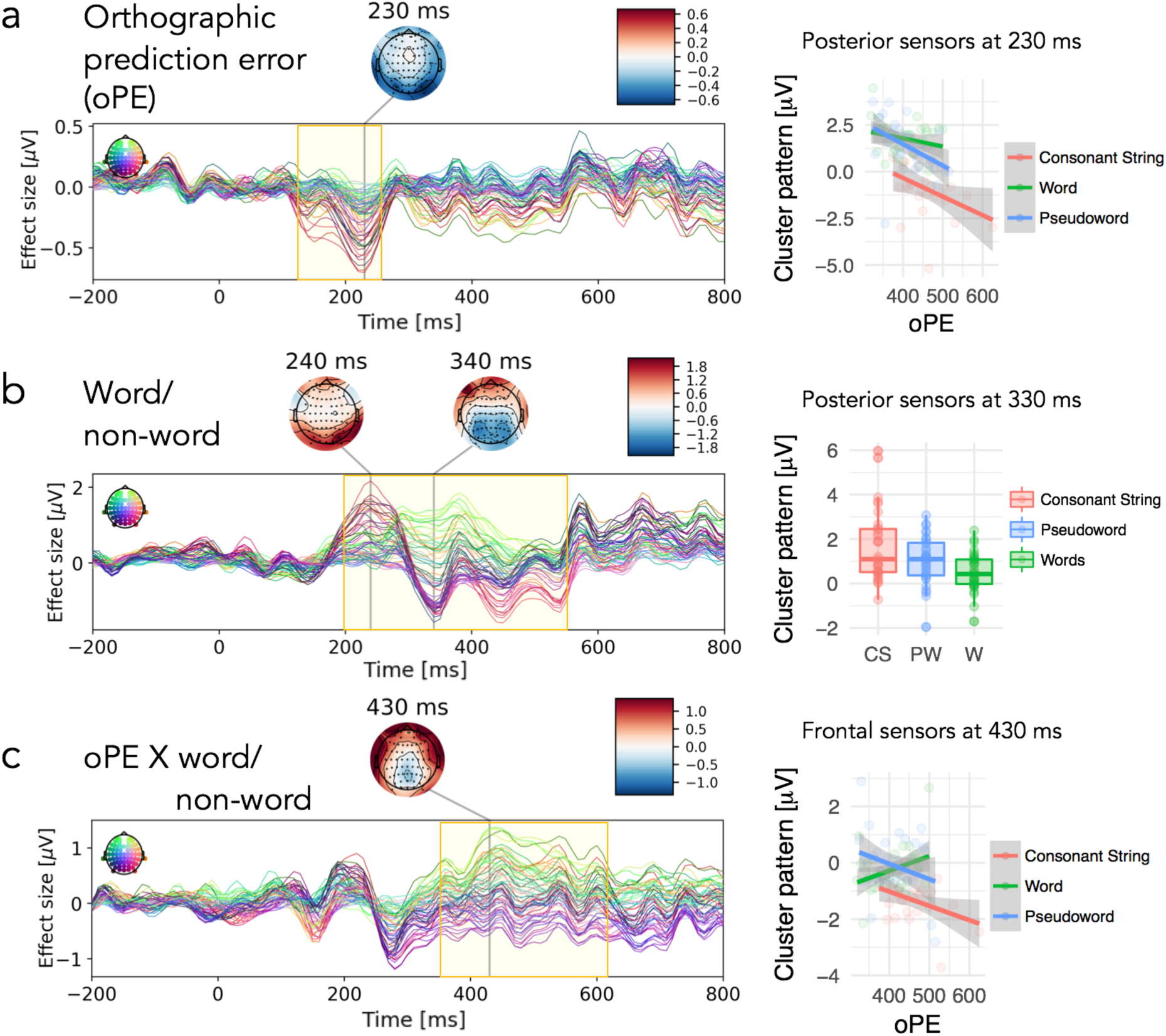
EEG results for silent reading of 200 words and 200 non-words (100 pronounceable pseudowords, 100 consonant strings): Timing of orthographic prediction error effects. Effect sizes from regression ERPs are presented as time courses for each sensor and time-point (left column; color coding reflects scalp position) with yellow areas marking time windows with significant activation clusters (see Supplemental Figure S3 for a more detailed visualization of the significance of spatio-temporal activation clusters). ERP results are shown for (a) the main effect of the orthographic prediction error (oPE), (b) the word/non-word effect, and (c) the oPE by word/non-word interaction. Results indicate significant oPE, word/non-word, and oPE by word/non-word effects starting around, 150, 200, and 360 ms, respectively. The right panel shows the activation patterns related to the significant activation clusters in more detail. Dots represent mean predicted μV across (a,c) all participants and items separated by oPE and stimulus category, and (b) all items separated by stimulus category, excluding confounding effects (see Materials and Methods). No significant activation clusters were found for the parameter representing the number of pixels. Boxplots represent the median (line), the data from the first to the third quartile (box), and ±1.5 times the interquartile range (i.e., quartile tree minus quartile one; whiskers). The frontal cluster includes the following sensors: AF3, AF4, AF7, AF8, F1, F2, F3, F4, F5, F6, F7, F8, SO1, SO2, FP1, FP2, Fz. The posterior cluster includes the following sensors: O2, O1, Oz, PO10, PO3, PO4, PO7, PO8, PO9, POz.

As in the fMRI study, we found no effect of the bottom-up input as such (pixel count), even though it is well-established that manipulations of physical input contrast (as determined, e.g., by the strength of luminance Johannes, Münte, Heinze, & Mangun, 1995) can increase the amplitude of early ERP components starting at around 100 ms. We performed an explicit model comparison between statistical models, including the orthographic prediction error compared to a model including the number of pixels parameter (analogous to the analysis of behavioral data), for both time windows in which the orthographic prediction error was relevant (early fixed effect and later interaction). In both time windows the model including the orthographic prediction error resulted in better fit (AIC difference: 16 at 230 ms at posterior sensors: χ^2^(0) = 16.0; p < .001; and 5 at 430 ms at frontal sensors: χ^2^(0) = 5.0; p < .001). Even when investigating the combined models, including both parameters, we found a tendency for a better fit in the oPE only model (AIC difference: 3 at 230 ms at posterior sensors and 3 at 430 ms at frontal sensors).

To summarize, EEG results converge with behavioral and fMRI results. They suggest that relatively early on in the cortical visual-perceptual processing cascade, the amount of perceptual processing devoted to the orthographic percept is smallest for letter-strings with highly expected visual features (i.e., low orthographic prediction error). 100 to 200 ms later, i.e., in a time window strongly associated with semantic processing (Kutas & Federmeier, 2011), the prediction error effect was selectively reversed for words, and thus started to differentiate between the two stimulus categories. This finding mirrors behavioral results and activation patterns in the anterior temporal lobe and prefrontal cortex found in the fMRI dataset. In sum, these results support the PEMoR’s proposal that orthographic representations are optimized early during visual word recognition, and that the resulting orthographic prediction error is the basis for subsequent stages of word recognition.

#### Applying the Prediction Error Model of Reading to handwritten script

The electronic fonts used for all the above-reported experiments introduce a highly regular structure that favors some of the PEMoR’s core processes, like the calculation of the orthographic prediction error (i.e., the prior). We showed above that when reducing the high regularity of computerized script by visual noise, reading performance decreases, and the orthographic prediction error becomes less relevant for describing reading behavior. To demonstrate that the PEMoR may be applicable also in less constrained settings with less regular visual-orthographic input, we applied a variant of this model to account for the ‘readability’ of naturalistic handwritings obtained from 10 different writers. The extreme variability of different handwritings strongly influences their readability (compare Figs. 6a,b). The visual-orthographic predictions which have here (for pragmatic reasons; see Materials and Methods) been implemented based on single letters and separately for each handwriting, accordingly vary substantially in strength and precision between individual handwritings (cp. ‘prediction’ of Fig. 6a,b). We define ‘prediction strength’ in terms of the summed darkness of all gray values of the prediction image, i.e., the mean gray value across pixels. We observe that ‘stronger’ priors are associated with lower oPEs across handwritings (Fig. 6c; linear mixed model statistics: Estimate: −0.05; SE = 0.01; t = 7.4), whereby calculating two oPE values per writer (i.e., one for upper and one for lower case letters). Here, we implemented the oPE calculation by subtracting the respective writer’s handwriting prediction, i.e., the mean across all letter images, from each letter sample provided by the same writer. Again the prediction errors are the sum of all grayscale values from the resulting difference images. In the final step, we estimated the mean prediction error for upper and lower case letters, respectively. Note that the oPE estimate for handwritings was normalized (i.e., divided) by the number of black/gray pixels of the letter image since this number differed drastically between scripts (e.g., compare Examples in Fig. 6a and b).

**Figure 6.**
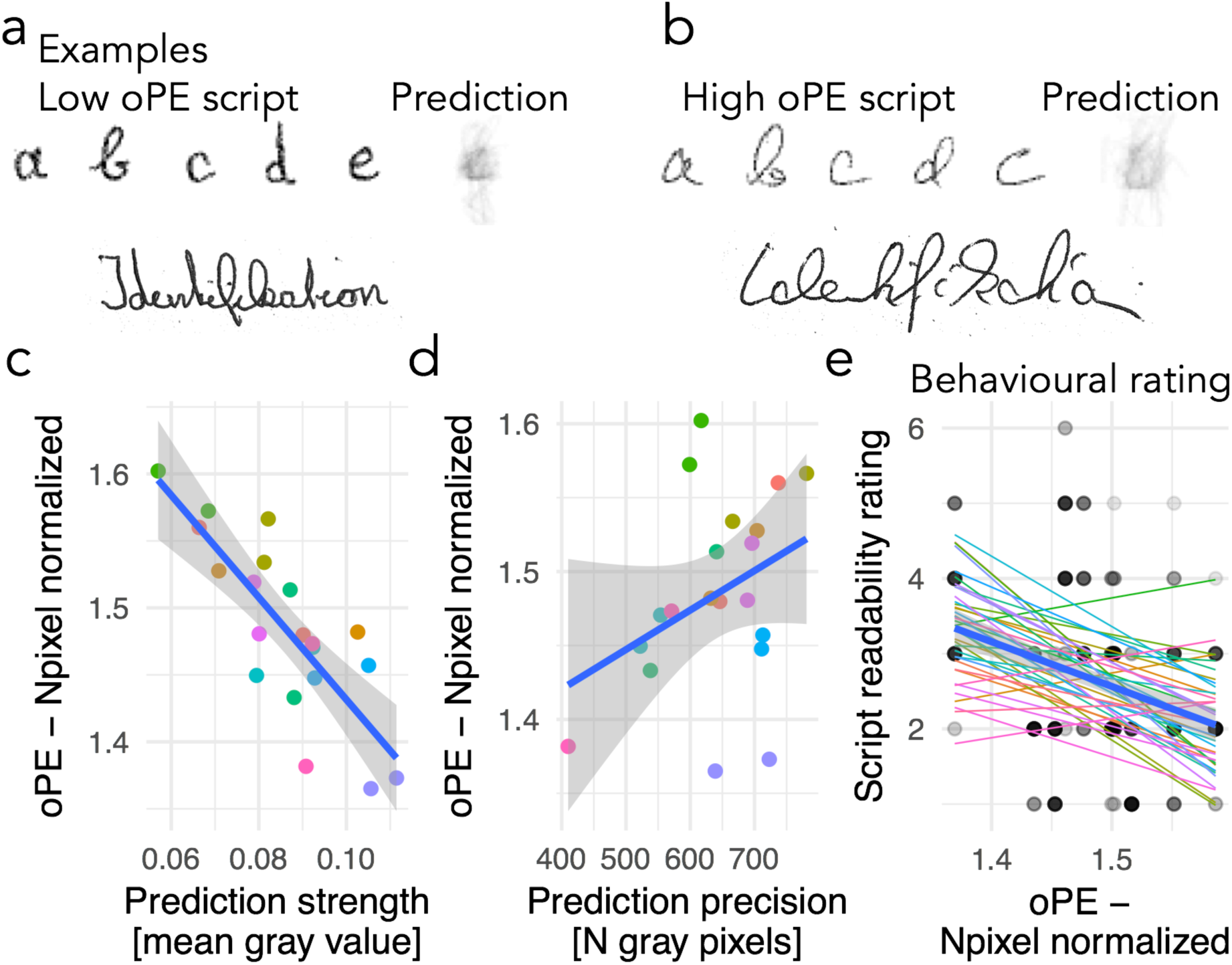
Applying the PEMoR to the perception of handwriting. (a,b) Examples for two (out of 10) empirically obtained handwritings. Writers provided samples of single letters, and a (letter-level) orthographic prediction was estimated based on all 26 letters (here demonstrated for lower-case letters). In addition, the word *Identifikation* (identification) is presented for both handwritings, as an example (out of 10 words obtained). These words were used to acquire the readability ratings from independent subjects. (c) Relationship between prediction strength and the mean orthographic prediction error across all letters for each script. Note that the oPE estimate for handwritings was normalized by the number of black/gray pixels of the letter image since this number differed drastically between scripts (e.g., compare Examples in a and b). This result is based on a linear mixed model with 20 observations, i.e., an oPE and a prediction strength estimate each for lower case and for upper case letters for each of the 10 writers. (d) Relationship between the precision of the prediction and the orthographic prediction error. The color of the dots reflects each of 10 individual scripts, separately for upper- and lower-case letters. (e) Readability ratings for each script in relation to the orthographic prediction error (combined across lower- and upper-case letters). The blue line reflects the overall relationship, while thin lines represent the 38 raters.

The ‘precision’ of the prediction can be represented by the inverse of the number of gray pixels included in the prediction image; more precise predictions are more focused and less distributed, and also elicit lower orthographic prediction errors (Fig. 6d; 0.02; SE = 0.01; t = 2.1; see Supplemental Table 1 for full results). Finally, we obtained the rated readability of each handwriting based on ten written words and observed that the readability is higher for handwritings that produce lower prediction errors (Fig. 6e; 38 raters; Estimate: −5.9; SE = 1.0; t = 6.2). These results demonstrate that (variants of) the PEMoR can account for reading processes not only in highly formalized stimuli but also in more naturalistic settings.

## Discussion

Here we investigated if an efficient neuronal code representing the visual input signal of words, i.e., analogous to the neural representation underlying the end stopping phenomenon for oriented lines, is the basis of proficient reading. We found that our Prediction Error Model of Reading (PEMoR) is a plausible account. Unexpectedly, since the focus of the PEMoR is visual processing, the resulting prediction error, i.e., the non-redundant and thus informative part of the visual percept, represents not only the visual but also orthographic word information. We concluded this from the correlations between various lexicon-based descriptors of words that are associated with orthographic stages of visual word recognition, and the prediction error representation. Our empirical results also support this conclusion: We found that the orthographic prediction error (i) accounts for word identification behavior, (ii) explains brain activation in visual-perceptual systems of the occipital cortex, (iii) explains brain activation as early as 150 ms after the onset of the letter-string, and (iv) contributes to high-level lexical processing that underlies word recognition behavior. We inferred the latter from the finding that both brain activation (i.e., BOLD activation in frontal areas and the N400 ERP effect) and lexical decision behavior showed comparable interaction effects between lexicality status (word/non-word) and the orthographic prediction error. Also, the PEMoR provides a quantitative estimate of the reduction of the full bottom-up signal achieved by the subtraction of the word-knowledge based prediction (i.e., in our data between 31 and 37% on average depending on language, with an upper limit of 51% at the level of the individual word). Finally, we have provided initial evidence that the principles of predictive coding may also apply to more naturalistic reading situations, by demonstrating that the orthographic prediction error can account for individual differences in the readability of handwriting. In sum, our findings indicate that the basis for fast access to the meaning of written words is an optimized neuronal representation that highlights unpredictable visual-orthographic word information.

We also found evidence that the reliance on the orthographic prediction error in word recognition is related to the perceptual quality of the stimulus. We showed that in case the visual occurrence of the stimulus is less predictable, e.g., due to visual noise, behavioral performance in the lexical decision task is better accounted for by our approximation of the original bottom-up input (i.e., the pixel count; see above) than by the optimized prediction error based signal. As described previously (Rao & Ballard, 1999), efficient coding in a predictive system relies on the structure present in the stimulus. If the structure is compromised, as is the case when adding visual noise, the predictive system breaks down as predictions become weaker and less precise. As a consequence, word processing relies on less efficient neuronal codes under such conditions.

Most results reported here relied on experiments with fixed word lengths, while naturalistic reading involves considerably more variability at the level of the input. However, para-foveal vision provides information about word length to the visual system before the actual processing of the word (Schotter et al., 2012) so that it is, in fact, often possible to dynamically implement best-fitting visual-orthographic predictions (priors) online during reading. This would, in principle, allow for optimized sensory processing as described by the PEMoR in natural reading situations. While this specific hypothesis must be tested in future studies, it fits well with previous theoretical proposals which have acknowledged the integration of top-down predictions from multiple linguistic domains (for example at the phonological, semantic, or syntactic level; DeLong et al., 2005; Eisenhauer, Fiebach, & Gagl, 2019; Nieuwland et al., 2018; Price & Devlin, 2011). Critically, our results go beyond these earlier models by demonstrating that top-down guided expectations are implemented already onto early cortical stages of visual-orthographic processing.

The so-far dominant model of visual word recognition in the brain (Dehaene & Cohen, 2011; Dehaene et al., 2005) postulates that words are ‘assembled’ bottom-up along the visual pathway, starting with symbolic representations of letter features (i.e., oriented lines) up to successively more complex higher-order representations. In this and similar models (including computational accounts of visual word recognition, e.g., Coltheart et al., 2001), the bottom-up ‘assembly’ of a letter or word representations is based on the full visual input signal. We have shown (in Fig. 2d) that the representational similarity space defined by the images of the word stimuli used in this study (i.e., the visual input) and the similarity space defined by the optimized orthographic prediction error images correlate to a high degree. This finding indicates that the PEMoR computation preserves the critical information for word identification in the orthographic prediction error representations. Consequently, we assume that there is a possibility that letter and word representations are assembled, as assumed in the currently dominant models, based on the prediction error representation. The Prediction Error Model of Reading, thus, does, in principle, not contradict other current models of visual word recognition. In contrast, PEMoR specifies explicitly and in a testable manner the nature of cortical representations that are generated early (i.e., prior to lexical access) in the perceptual process and on which all subsequent processing stages operate. Prediction-based top-down optimization of the visual-orthographic input, as proposed here, offers a possible specification of a previously underspecified aspect of visual word recognition models, i.e., the nature of representations that are active between visual perception and more abstract processes of lexical access.

Predictive coding-based theories, in general, assume that ‘higher-level’ stages of cortical processing aim at prediction error minimization as a core computation (Friston, 2005; Price & Devlin, 2011). As described in the Introduction section, such an account would be fundamentally different from most assumptions in established models of visual word recognition. Given the successful empirical evaluations reported in the present study, a future challenge for model development should be to incorporate an orthographic prediction error signal into current model assumptions about orthographic processing. For example, when conceptualizing word recognition as a process of evidence accumulation process with the goal of identifying the respective word in the mental lexicon (similar as previously described in Engbert et al., 2005; Gagl, Richlan, Ludersdorfer, Sassenhagen, & Fiebach, 2016; Ratcliff, Gomez, & McKoon, 2004; Summerfield & de Lange, 2014), the orthographic prediction error most likely serves as only one among several sources of evidence, particularly during natural reading which only rarely involves the processing of isolated words. Almost inevitably, contextual constraints will be operative. At the semantic level, it is well established that word-elicited brain activation (like the N400 component of the EEG or MEG) decreases if the word is predictable from its semantic context (e.g., Dambacher, Kliegl, Hofmann, & Jacobs, 2006; Dimigen, Sommer, Hohlfeld, Jacobs, & Kliegl, 2011; Eisenhauer et al., 2019; see also Kutas & Federmeier, 2011 for a review). Similarly, eye fixation durations during natural reading are reduced with increasing context-dependent word predictability (e.g., Hawelka, Schuster, Gagl, & Hutzler, 2015; Kliegl, Grabner, Rolfs, & Engbert, 2004; Kliegl et al., 2006). Syntactic-level predictions, in contrast, were identified as small in recent investigations (Nicenboim, Vasishth, & Rösler, 2019; Nieuwland et al., 2018). Beyond contextual constraints, it is also plausible that phonological information (as proposed by Price & Devlin, 2011 or morphological constraints may be applied in a top-down manner onto bottom-up visual-orthographic processing. Combined, these considerations suggest that all relevant levels of linguistic representation involved in language comprehension are likely to interact pro-actively, if possible, during the identification of each upcoming word. As a result, top-down predictions about expected upcoming words may in naturalistic situations often be much more specific than lexical knowledge-based priors investigated here, as they are based on candidate sets constrained by multiple sources of linguistic expectations. The work presented here, thus, should be understood as proof of principle under highly unconstrained conditions, i.e., the recognition of single words in isolation.

In this study, we presented evidence for a quantitative implementation of orthographic prediction error representations with stimuli (i) in Courier new, i.e., a monospaced font, (ii) for the effect of degraded stimuli on the effect of the orthographic prediction error, and (iii) for a possible application to more naturalistic conditions by using handwritten stimuli. One possible limitation of the current study is that we did not include a systematic investigation of different fonts (including proportional fonts). However, variation in font type is one among several sources of visual noise in the stimuli, and our investigation of how visual noise influences the reliance of word recognition processes on the prediction error signal provides a first hint of how font variation (as well as variations in size, position, etc.) may influence prediction-based word recognition processes. A systematic investigation of font effects, however, is an important future step of further exploring the here-proposed model. For example, it is of interest if and how readers adapt dynamically to new fonts, which might allow providing recommendations to inform design decisions for written outlets (e.g., for newspapers, or webpages) on the costs of using multiple fonts (e.g., see Gagl, 2016 for a recommendation concerning font color).

In sum, we demonstrate that during reading, the visual input signal is optimized by ‘explaining away’ redundant parts of the visual input based on top-down orthographic predictions. This study, accordingly, provides strong evidence that reading follows domain-general mechanisms of predictive coding during perception (Clark, 2013) and is also consistent with the influential hypothesis of a Bayesian brain, which during perception continuously combines prior knowledge and new sensory evidence (Friston, 2005; Knill & Pouget, 2004). We propose that the result of this optimization step, i.e., an orthographic prediction error signal, is the efficient neuronal code on which subsequent, ‘higher’ levels of word recognition operate, including the activation of word meaning. These data provide the basis for a new understanding of early, i.e., pre-lexical orthographic stages of visual word recognition, rooted in a strong and widely accepted, domain-general neurophysiological model – prediction-based perception (Friston, 2005; Rao & Ballard, 1999). At the same time, our results provide crucial converging evidence in support of predictive coding theory.

## Acknowledgments

We thank Rebekka Tenderra, Anne Hoffmann, Jan Jürges, and Kirsten Hilger for help with EEG data acquisition. In addition, we thank Mark D’Esposito, David Poeppel, Matt Davis, and Ulrike Basten for helpful comments on a previous version of the manuscript. The research leading to these results has received funding from the European Community’s Seventh Framework Programme (FP7/2013) under grant agreement n° 617891 awarded to CJF and from the European Community’s Horizon 2020 Programme (2016) under grant agreement n° 707932 awarded to BG.

## Author contributions

B.G. and C.F. wrote and revised the manuscript and all of the authors edited the final drafts. B.G., J.S., and C.F. conceptualized the model. B.G. and S.H. implemented the model. B.G. implemented the model simulations, conceptualized behavioral experiments, and behavioral data analysis. B.G. and K.G. implemented behavioral data acquisition, and the handwriting adaptation. B.G., J.S., and S.H. designed, measured and analyzed the EEG study. B.G. and F.R. designed, measured and analyzed the fMRI study.

## Data and materials availability

Data and analysis scripts are available at https://osf.io/d8yjc/

## Supplementary Materials

**Figure S1.**
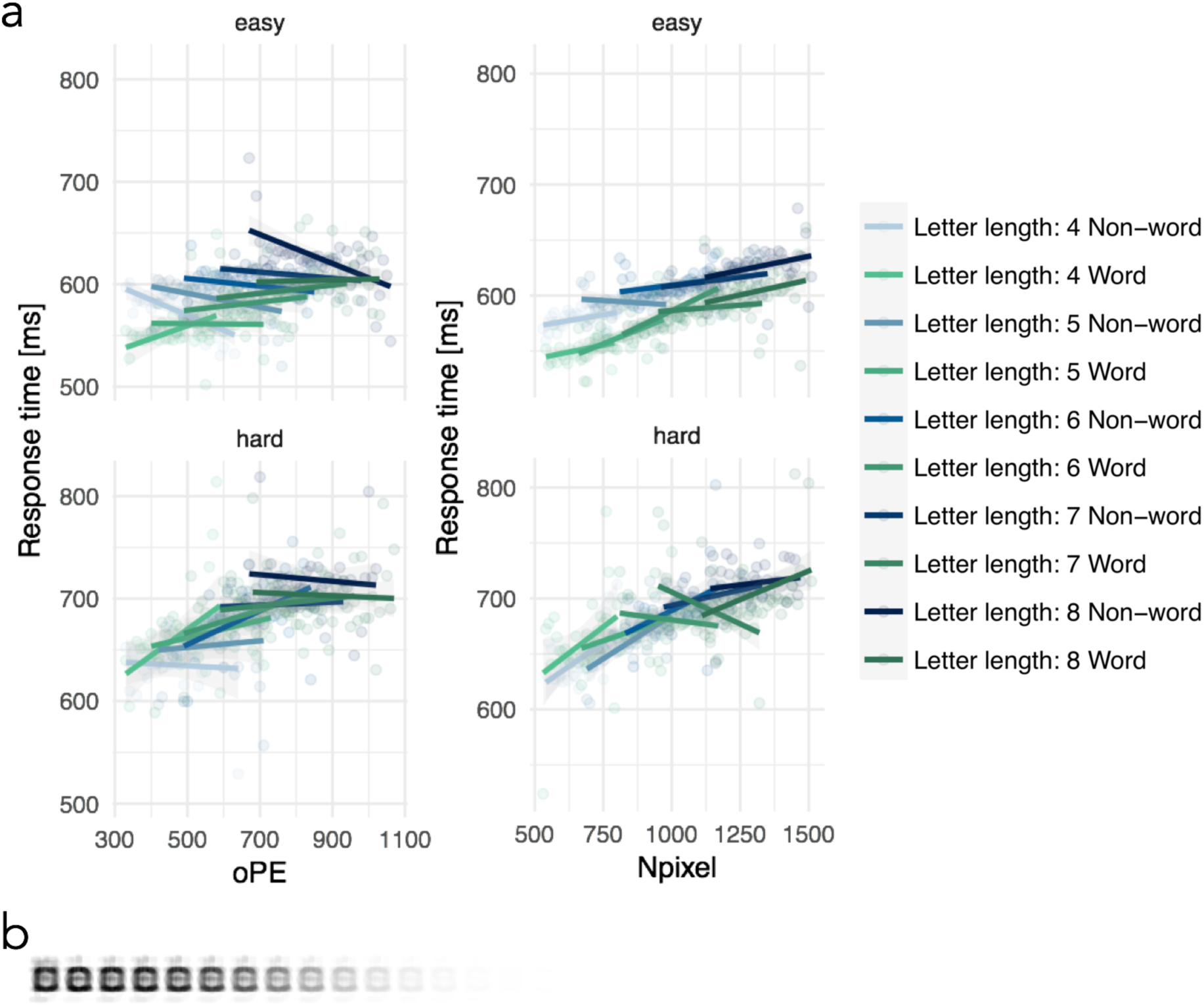
Behavioral evaluation including multiple word lengths. (a) Response times aggregated across participants from the British lexicon (BLP) project (Keuleers et al., 2012) for the word lengths 4-8. The left panel shows the word/non-word by orthographic prediction error (oPE) interaction and the right panel shows the word/non-word by number of pixels (Npixel) interaction for each word length separately. In addition, the upper panel shows letter strings that are correctly categorized in nearly all cases (accuracy > .95) and the lower panel shows the response times to the items, which were less accurately processed (i.e., accuracy < .95). The median split resulted in a subset of the BLP (i.e., the easy words) which are roughly comparable to words used in the previous experiments (e.g. see Fig. 3), as the BLP study includes a large number of very rare words (median log. word frequency per million is .3). Bluish colors represent non-words (N) and greenish colors represent words (W), while the hue of the colors reflects word length (i.e., bright to dark reflects short to long letter strings). For both effects, we first estimated linear regression models with either the oPE or the Npixel effect and allowing interactions with word/non-word status, word length, and accuracy. Note that the oPE in this first analysis was based on length-specific predictions (i.e., for the estimation of the oPE of four-letter words, all four-letter words of the lexicon were included in the prediction). For the oPE model, a significant four-way interaction was found (estimate = −1.078e-04; SE = 4.199e-05; *t* = −2.567). Separating hard vs. easy words allowed us to disentangle the four-way interaction: In easy words/non-words, we found a consistent (i.e., across length levels) oPE by word/non-word interaction (estimate = 1.530e-04; SE = 4.047e-05; *t* = 3.780) in the same direction as previously shown (positive effect for words and a negative effect for non-words). For hard words/non-words, we found that the oPE by word/non-word interaction was inconsistent across letter length levels, which was indicated by a significant oPE and letter length interaction (estimate = −3.530e-05; SE = 8.092e-06; *t* = −4.363). In addition, for the hard words both the oPE by word/non-word interaction (estimate = −1.685e-04; SE = 6.905e-05; *t* = −2.440) and the main effect of oPE were reversed (estimate = 2.828e-04; SE = 5.802e-05; *t* = 4.874 compare to estimate = −1.000e-04; SE = 2.440e-05; *t* = −4.101, for easy words). For the Npixel model, no four-way interaction and no Npixel interaction or main effect were found. In sum, in this analysis we showed that the oPE by word/non-word interaction shown previously for word lengths of five letters (see main text) is consistent for easy-to-process English items with word lengths from 4-8 letters. Secondly, the word/non-word by orthographic prediction error interaction was also reliable when the prediction included all words of all letter lengths from the English lexicon (see part b of this Figure) and the orthographic prediction error estimation was based on this length-unspecific prediction (estimate: 0.02; SE=0.007; t=3.349). (b) Letter-length unspecific prediction for English based on ∼60,000 English words from the SUBTLEX database (Heuven et al., 2014).

**Figure S2.**
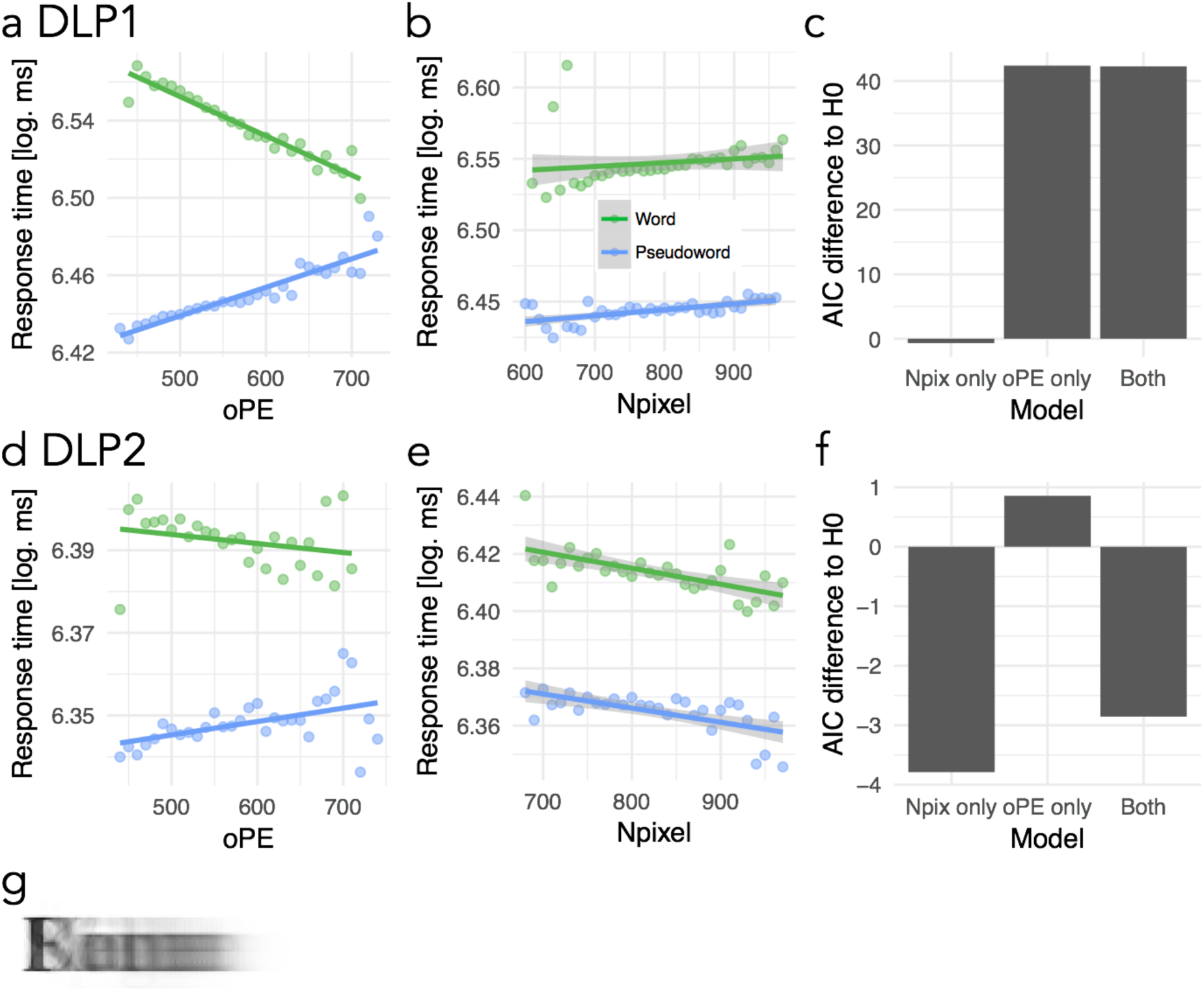
Dutch lexical decision behavior and prediction using a proportional script. (a) Effect of the orthographic prediction error parameter, (b) number of pixels parameter and (c) showing the same model comparisons as implemented in Figure 3 for the data from the first Dutch lexicon project (DLP1; (Keuleers, Diependaele, & Brysbaert, 2010); 4,305 five-letter stimuli; 39 participants) and the same effects and model comparisons for the second Dutch lexicon project (DLP2; (Brysbaert, Stevens, Mandera, & Keuleers, 2016); 3,145 five-letter stimuli; 81 participants) are presented in (d,e,f). Before going into the details of the two studies one has to note that the patterns we have found in the data in relation to our parameters of interest do not replicated within these two Dutch studies and, in addition, do not replicate with the findings from German, English, and French shown in Figure 3. In general, this is difficult for the interpretations of the results. For the DLP1 pattern we found a significant interaction of the orthographic prediction error with word/non-words and no significant effect of number of pixels. The interaction pattern in contrast to the findings in other languages (Fig. 3a), however, was qualitatively different as it showed a negative orthographic prediction error effect for words and a positive effect for non-words. The pattern is exactly the inverse from all other languages. Still model comparisons highlighted that the orthographic prediction error was relevant for the model fit since the predictor increased the model fit with no further increase of fit when the number of pixel parameter was included. None of these findings could be replicated in the DLP2 dataset, showing no significant fixed effects or interactions and no substantial changes in model fit relation to the null model. (g) Prediction image from a PEMoR implementation using five-letter words with a proportional Times New Roman script.

**Figure S3.**
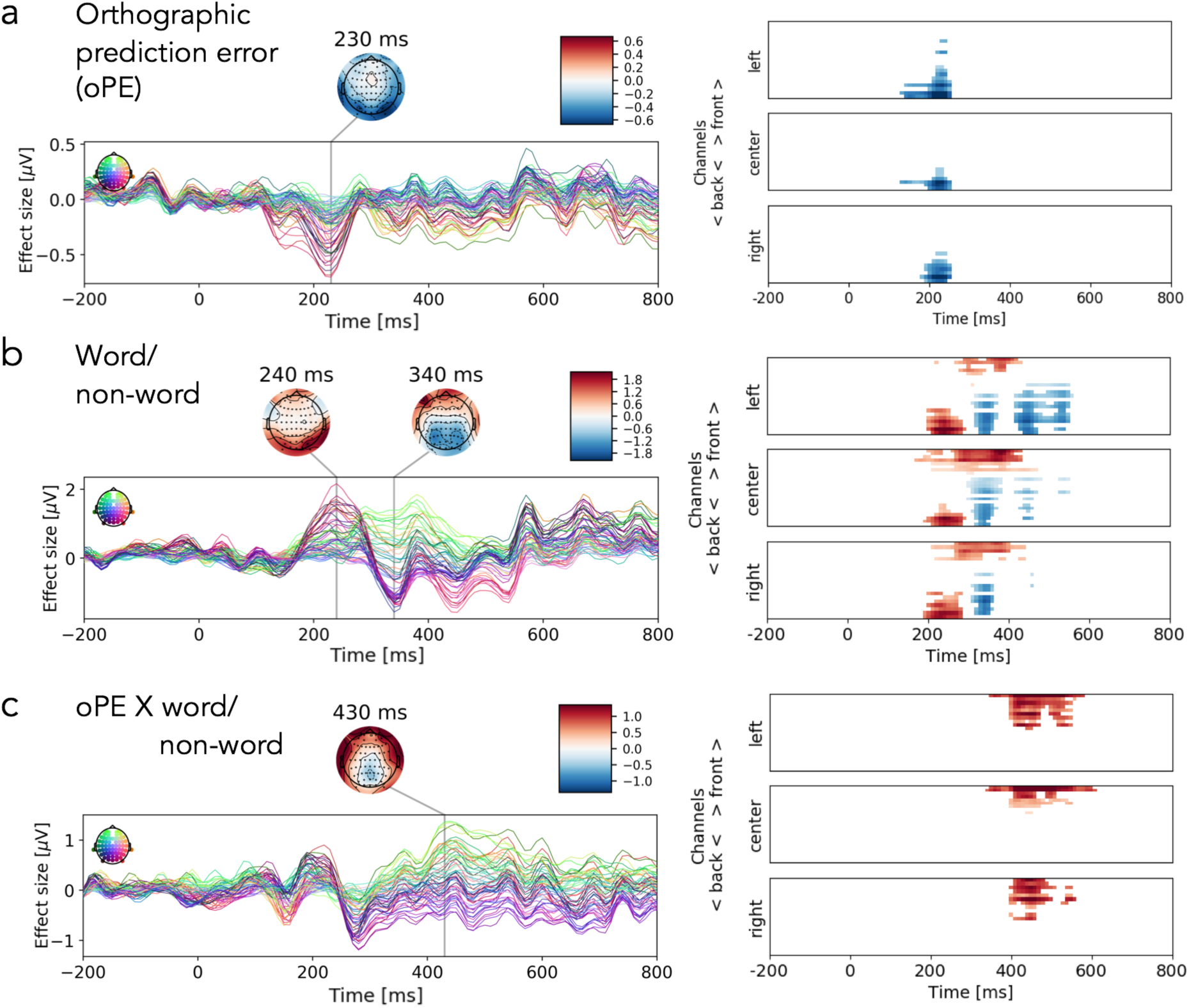
Detailed description of significant activation clusters in the EEG study for (a) the orthographic prediction error; (b) word/non-word effect; (c) interaction of word/non-word and the orthographic prediction error. On the left, the effect sizes from regression ERPs are presented as time courses for each sensor and time-point (color coding reflects scalp position). This part of the Figure reproduces Figure 5. The right column displays time courses with one line per channel, masked by significance using cluster statistics (see Methods for details; Maris & Oostenveld, 2007).

**Table S1.**
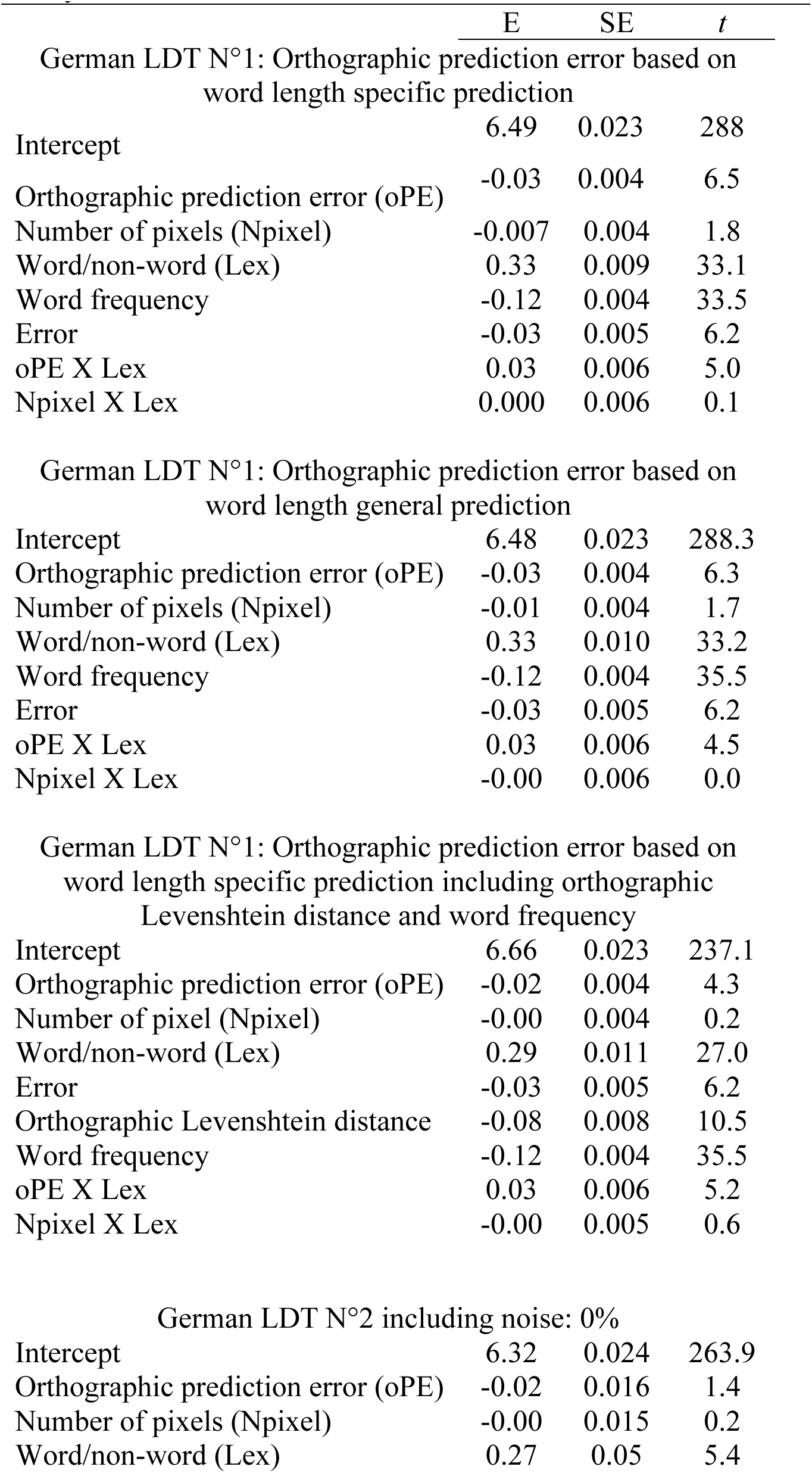
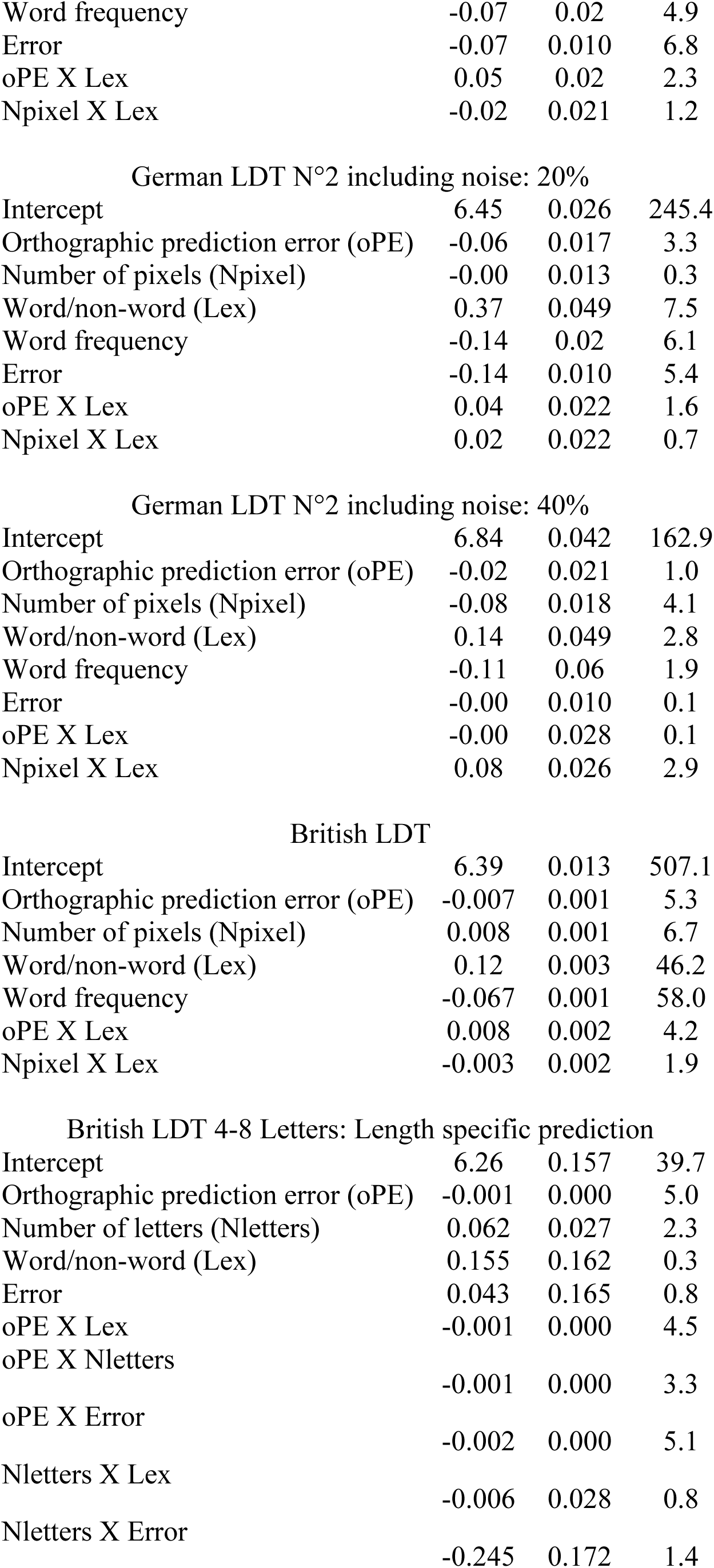
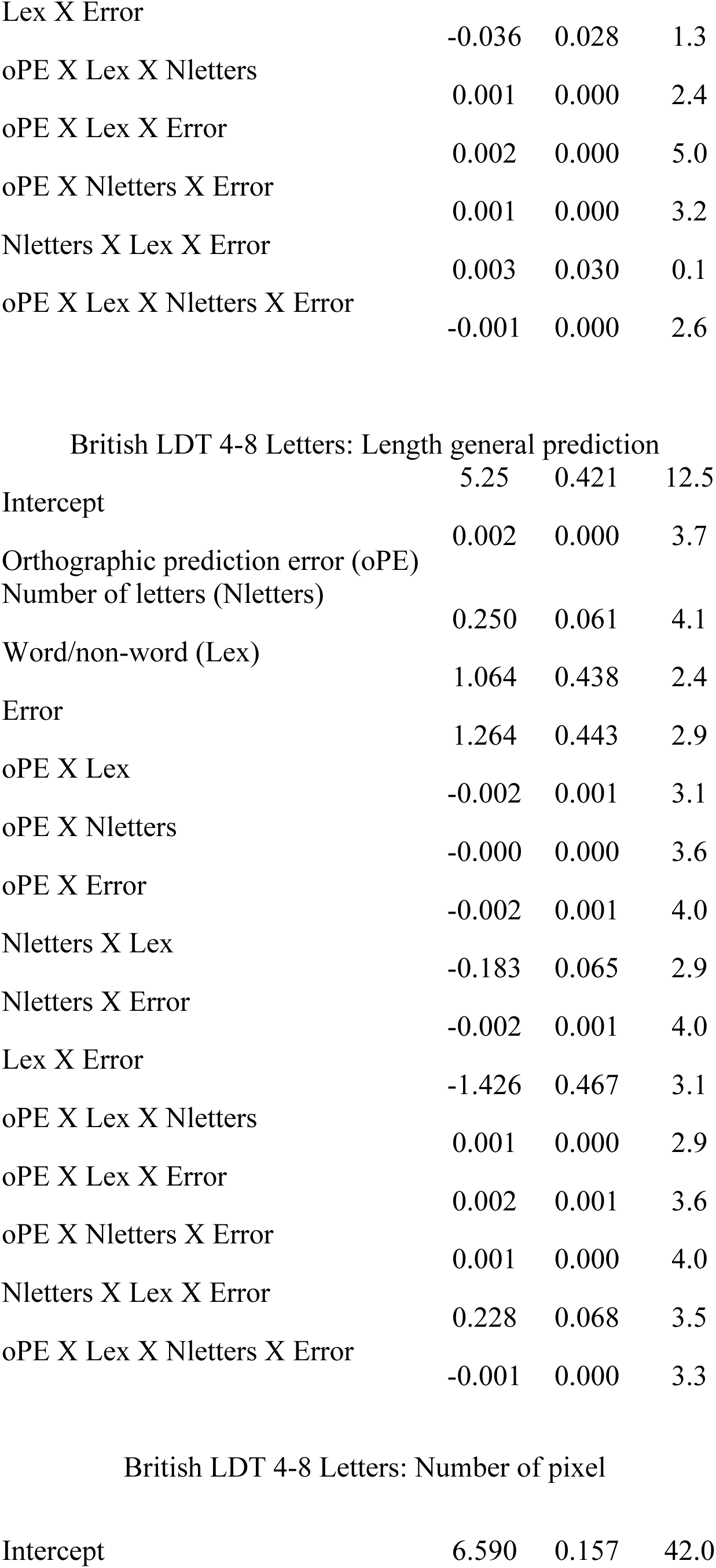
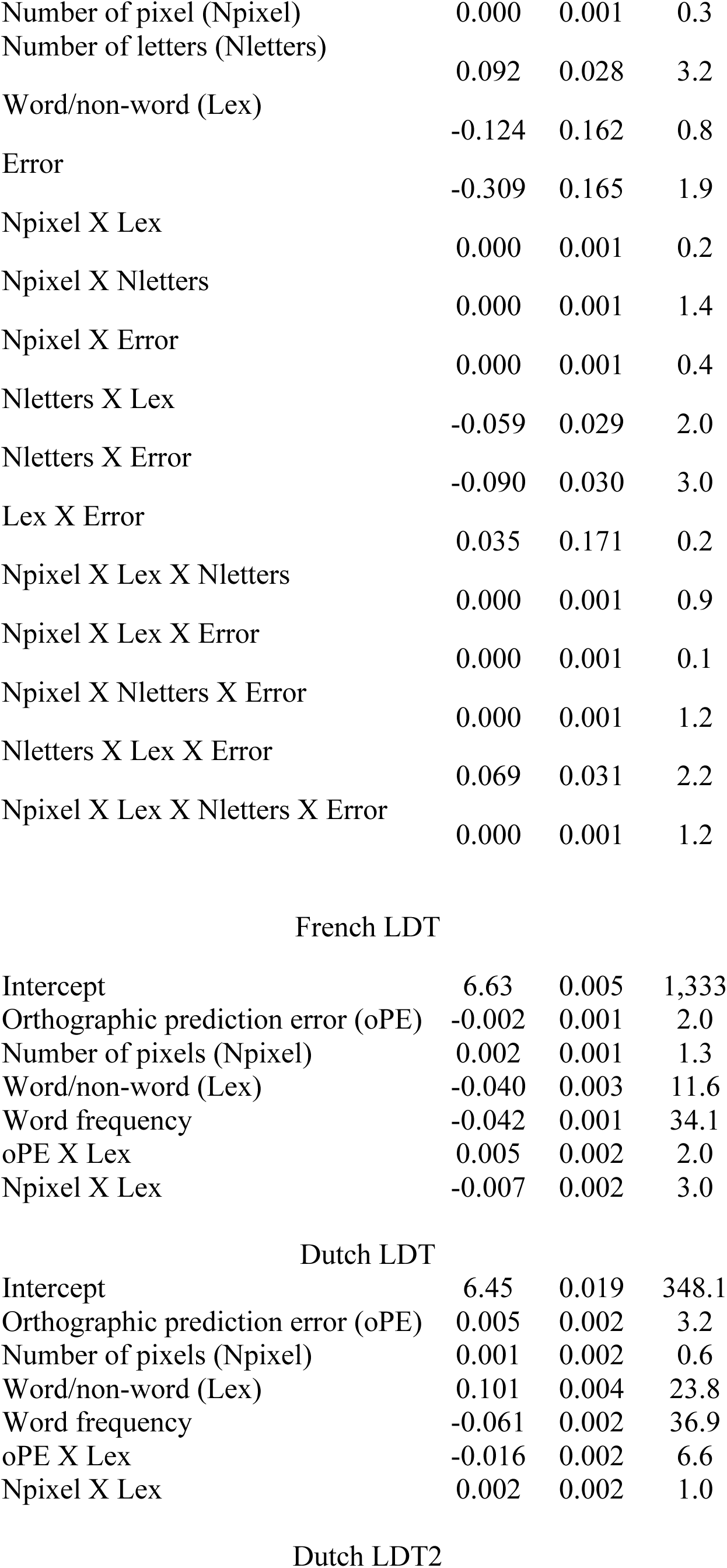
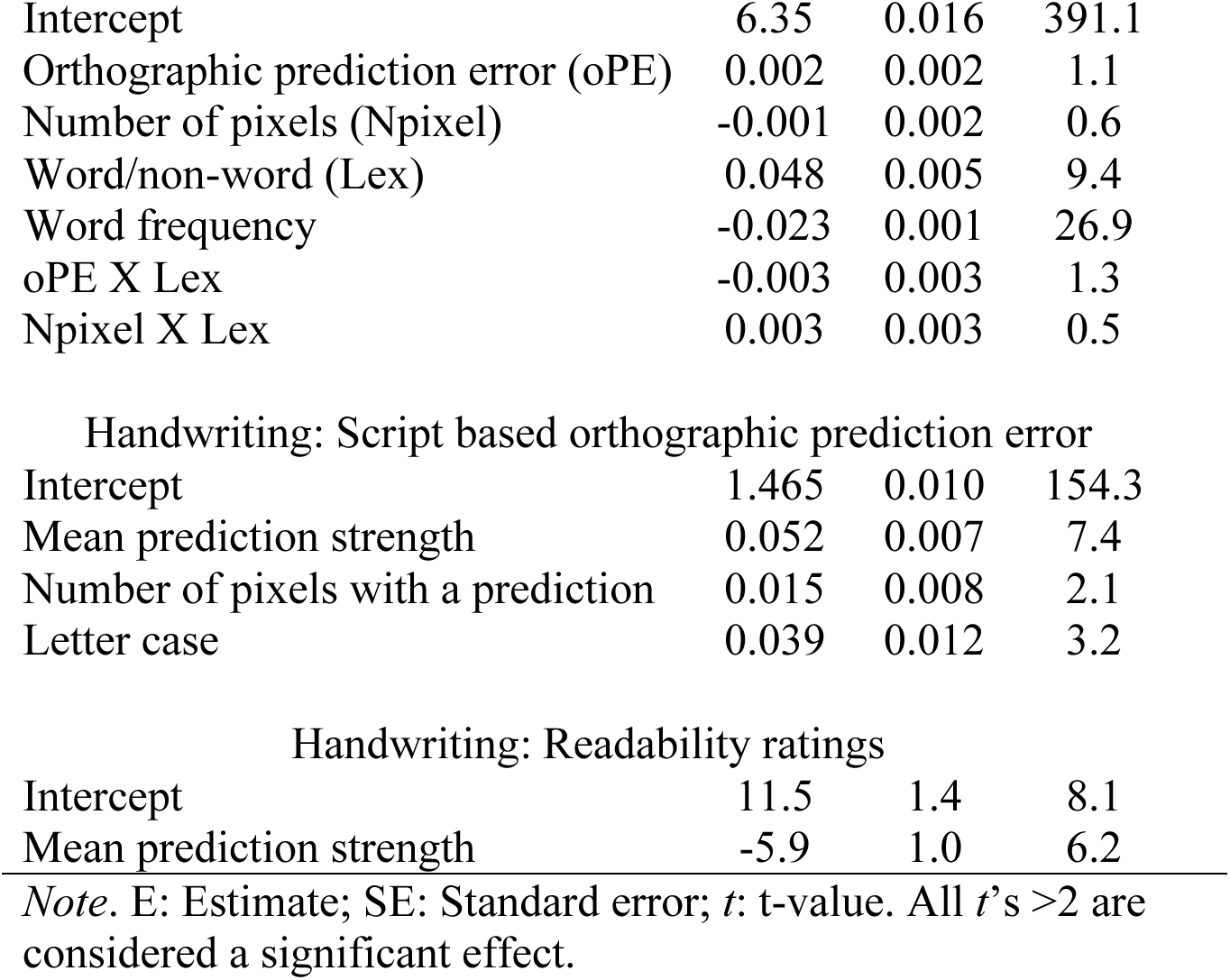
Results from linear mixed model regression analysis (with the exception of the British data including multiple word lengths was estimated based on word aggregated data) for the behavioral lexical decision tasks (LDT) and handwriting analyses.

|  | E | SE | *t* |
| --- | --- | --- | --- |
| German LDT N°1: Orthographic prediction error based on word length specific prediction |  |  |  |
| Intercept | 6.49 | 0.023 | 288 |
| Orthographic prediction error (oPE) | -0.03 | 0.004 | 6.5 |
| Number of pixels (Npixel) | -0.007 | 0.004 | 1.8 |
| Word/non-word (Lex) | 0.33 | 0.009 | 33.1 |
| Word frequency | -0.12 | 0.004 | 33.5 |
| Error | -0.03 | 0.005 | 6.2 |
| oPE X Lex | 0.03 | 0.006 | 5.0 |
| Npixel X Lex | 0.000 | 0.006 | 0.1 |
| German LDT N°1: Orthographic prediction error based on word length general prediction |  |  |  |
| Intercept | 6.48 | 0.023 | 288.3 |
| Orthographic prediction error (oPE) | -0.03 | 0.004 | 6.3 |
| Number of pixels (Npixel) | -0.01 | 0.004 | 1.7 |
| Word/non-word (Lex) | 0.33 | 0.010 | 33.2 |
| Word frequency | -0.12 | 0.004 | 35.5 |
| Error | -0.03 | 0.005 | 6.2 |
| oPE X Lex | 0.03 | 0.006 | 4.5 |
| Npixel X Lex | -0.00 | 0.006 | 0.0 |
| German LDT N°1: Orthographic prediction error based on word length specific prediction including orthographic Levenshtein distance and word frequency |  |  |  |
| Intercept | 6.66 | 0.023 | 237.1 |
| Orthographic prediction error (oPE) | -0.02 | 0.004 | 4.3 |
| Number of pixel (Npixel) | -0.00 | 0.004 | 0.2 |
| Word/non-word (Lex) | 0.29 | 0.011 | 27.0 |
| Error | -0.03 | 0.005 | 6.2 |
| Orthographic Levenshtein distance | -0.08 | 0.008 | 10.5 |
| Word frequency | -0.12 | 0.004 | 35.5 |
| oPE X Lex | 0.03 | 0.006 | 5.2 |
| Npixel X Lex | -0.00 | 0.005 | 0.6 |
| German LDT N°2 including noise: 0% |  |  |  |
| Intercept | 6.32 | 0.024 | 263.9 |
| Orthographic prediction error (oPE) | -0.02 | 0.016 | 1.4 |
| Number of pixels (Npixel) | -0.00 | 0.015 | 0.2 |
| Word/non-word (Lex) | 0.27 | 0.05 | 5.4 |
| Word frequency | -0.07 | 0.02 | 4.9 |
| Error | -0.07 | 0.010 | 6.8 |
| oPE X Lex | 0.05 | 0.02 | 2.3 |
| Npixel X Lex | -0.02 | 0.021 | 1.2 |
| German LDT N°2 including noise: 20% |  |  |  |
| Intercept | 6.45 | 0.026 | 245.4 |
| Orthographic prediction error (oPE) | -0.06 | 0.017 | 3.3 |
| Number of pixels (Npixel) | -0.00 | 0.013 | 0.3 |
| Word/non-word (Lex) | 0.37 | 0.049 | 7.5 |
| Word frequency | -0.14 | 0.02 | 6.1 |
| Error | -0.14 | 0.010 | 5.4 |
| oPE X Lex | 0.04 | 0.022 | 1.6 |
| Npixel X Lex | 0.02 | 0.022 | 0.7 |
| German LDT N°2 including noise: 40% |  |  |  |
| Intercept | 6.84 | 0.042 | 162.9 |
| Orthographic prediction error (oPE) | -0.02 | 0.021 | 1.0 |
| Number of pixels (Npixel) | -0.08 | 0.018 | 4.1 |
| Word/non-word (Lex) | 0.14 | 0.049 | 2.8 |
| Word frequency | -0.11 | 0.06 | 1.9 |
| Error | -0.00 | 0.010 | 0.1 |
| oPE X Lex | -0.00 | 0.028 | 0.1 |
| Npixel X Lex | 0.08 | 0.026 | 2.9 |
| British LDT |  |  |  |
| Intercept | 6.39 | 0.013 | 507.1 |
| Orthographic prediction error (oPE) | -0.007 | 0.001 | 5.3 |
| Number of pixels (Npixel) | 0.008 | 0.001 | 6.7 |
| Word/non-word (Lex) | 0.12 | 0.003 | 46.2 |
| Word frequency | -0.067 | 0.001 | 58.0 |
| oPE X Lex | 0.008 | 0.002 | 4.2 |
| Npixel X Lex | -0.003 | 0.002 | 1.9 |
| British LDT 4-8 Letters: Length specific prediction |  |  |  |
| Intercept | 6.26 | 0.157 | 39.7 |
| Orthographic prediction error (oPE) | -0.001 | 0.000 | 5.0 |
| Number of letters (Nletters) | 0.062 | 0.027 | 2.3 |
| Word/non-word (Lex) | 0.155 | 0.162 | 0.3 |
| Error | 0.043 | 0.165 | 0.8 |
| oPE X Lex | -0.001 | 0.000 | 4.5 |
| oPE X Nletters | -0.001 | 0.000 | 3.3 |
| oPE X Error | -0.002 | 0.000 | 5.1 |
| Nletters X Lex | -0.006 | 0.028 | 0.8 |
| Nletters X Error | -0.245 | 0.172 | 1.4 |
| Lex X Error | -0.036 | 0.028 | 1.3 |
| oPE X Lex X Nletters | 0.001 | 0.000 | 2.4 |
| oPE X Lex X Error | 0.002 | 0.000 | 5.0 |
| oPE X Nletters X Error | 0.001 | 0.000 | 3.2 |
| Nletters X Lex X Error | 0.003 | 0.030 | 0.1 |
| oPE X Lex X Nletters X Error | -0.001 | 0.000 | 2.6 |
| British LDT 4-8 Letters: Length general prediction |  |  |  |
| Intercept | 5.25 | 0.421 | 12.5 |
| Orthographic prediction error (oPE) | 0.002 | 0.000 | 3.7 |
| Number of letters (Nletters) | 0.250 | 0.061 | 4.1 |
| Word/non-word (Lex) | 1.064 | 0.438 | 2.4 |
| Error | 1.264 | 0.443 | 2.9 |
| oPE X Lex | -0.002 | 0.001 | 3.1 |
| oPE X Nletters | -0.000 | 0.000 | 3.6 |
| oPE X Error | -0.002 | 0.001 | 4.0 |
| Nletters X Lex | -0.183 | 0.065 | 2.9 |
| Nletters X Error | -0.002 | 0.001 | 4.0 |
| Lex X Error | -1.426 | 0.467 | 3.1 |
| oPE X Lex X Nletters | 0.001 | 0.000 | 2.9 |
| oPE X Lex X Error | 0.002 | 0.001 | 3.6 |
| oPE X Nletters X Error | 0.001 | 0.000 | 4.0 |
| Nletters X Lex X Error | 0.228 | 0.068 | 3.5 |
| oPE X Lex X Nletters X Error | -0.001 | 0.000 | 3.3 |
| British LDT 4-8 Letters: Number of pixel |  |  |  |
| Intercept | 6.590 | 0.157 | 42.0 |
| Number of pixel (Npixel) | 0.000 | 0.001 | 0.3 |
| Number of letters (Nletters) | 0.092 | 0.028 | 3.2 |
| Word/non-word (Lex) | -0.124 | 0.162 | 0.8 |
| Error | -0.309 | 0.165 | 1.9 |
| Npixel X Lex | 0.000 | 0.001 | 0.2 |
| Npixel X Nletters | 0.000 | 0.001 | 1.4 |
| Npixel X Error | 0.000 | 0.001 | 0.4 |
| Nletters X Lex | -0.059 | 0.029 | 2.0 |
| Nletters X Error | -0.090 | 0.030 | 3.0 |
| Lex X Error | 0.035 | 0.171 | 0.2 |
| Npixel X Lex X Nletters | 0.000 | 0.001 | 0.9 |
| Npixel X Lex X Error | 0.000 | 0.001 | 0.1 |
| Npixel X Nletters X Error | 0.000 | 0.001 | 1.2 |
| Nletters X Lex X Error | 0.069 | 0.031 | 2.2 |
| Npixel X Lex X Nletters X Error | 0.000 | 0.001 | 1.2 |
| French LDT |  |  |  |
| Intercept | 6.63 | 0.005 | 1,333 |
| Orthographic prediction error (oPE) | -0.002 | 0.001 | 2.0 |
| Number of pixels (Npixel) | 0.002 | 0.001 | 1.3 |
| Word/non-word (Lex) | -0.040 | 0.003 | 11.6 |
| Word frequency | -0.042 | 0.001 | 34.1 |
| oPE X Lex | 0.005 | 0.002 | 2.0 |
| Npixel X Lex | -0.007 | 0.002 | 3.0 |
| Dutch LDT |  |  |  |
| Intercept | 6.45 | 0.019 | 348.1 |
| Orthographic prediction error (oPE) | 0.005 | 0.002 | 3.2 |
| Number of pixels (Npixel) | 0.001 | 0.002 | 0.6 |
| Word/non-word (Lex) | 0.101 | 0.004 | 23.8 |
| Word frequency | -0.061 | 0.002 | 36.9 |
| oPE X Lex | -0.016 | 0.002 | 6.6 |
| Npixel X Lex | 0.002 | 0.002 | 1.0 |
| Dutch LDT2 |  |  |  |
| Intercept | 6.35 | 0.016 | 391.1 |
| Orthographic prediction error (oPE) | 0.002 | 0.002 | 1.1 |
| Number of pixels (Npixel) | -0.001 | 0.002 | 0.6 |
| Word/non-word (Lex) | 0.048 | 0.005 | 9.4 |
| Word frequency | -0.023 | 0.001 | 26.9 |
| oPE X Lex | -0.003 | 0.003 | 1.3 |
| Npixel X Lex | 0.003 | 0.003 | 0.5 |
| Intercept | 1.465 | 0.010 | 154.3 |
| Mean prediction strength | 0.052 | 0.007 | 7.4 |
| Number of pixels with a prediction | 0.015 | 0.008 | 2.1 |
| Letter case | 0.039 | 0.012 | 3.2 |
| Handwriting: Readability ratings |  |  |  |
| Intercept | 11.5 | 1.4 | 8.1 |
| Mean prediction strength | -5.9 | 1.0 | 6.2 |

*Note.* E: Estimate; SE: Standard error; *t*: t-value. All *t*'s >2 are considered a significant effect.

Table S2. Reliable activation clusters from the fMRI evaluation with respective anatomical labels

